# Mapping the contents of consciousness during musical imagery

**DOI:** 10.1101/2020.11.20.391375

**Authors:** Mor Regev, Andrea R. Halpern, Adrian M. Owen, Aniruddh D. Patel, Robert J. Zatorre

**Author notes:** Corresponding author: Mor Regev.

## Abstract

Humans can internally represent auditory information without an external stimulus. When imagining music, how similar are unfolding neural representations to those during the original perceived experience? Can rhythmic motion influence the neural representation of music during imagery as during perception? Participants first memorized six one-minute-long instrumental musical pieces with high accuracy. Functional MRI data were collected during: 1) silent imagery of melodies to the beat of a visual metronome; 2) same but while tapping to the beat; and 3) passive listening. During imagery, inter-subject comparison showed that melody-specific temporal response patterns were reinstated in right associative auditory cortices. When tapping accompanied imagery, the melody-specific neural patterns were extended to associative cortices bilaterally. These results indicate that the specific contents of conscious experience are encoded similarly during imagery and perception in the dynamic activity of auditory cortices. Furthermore, rhythmic motion can enhance the reinstatement of neural patterns associated with the experience of complex sounds, in keeping with models of motor to sensory influences in auditory processing.

## Introduction

Mental imagery, the conscious representation of sensory information without direct external stimulus, is arguably one of the primary components of human cognition. It plays an important role in memory retrieval, future planning, decision making, creativity, and emotional regulation (Baddeley and Logie, 1992; Moulton and Kosslyn, 2009; Lucas et al., 2010; Keogh and Pearson, 2011; Palmiero et al., 2011; Amit and Greene, 2012; Schacter et al., 2012). In mental imagery, previously experienced sensory content occupies our conscious mind, reinstating old events or composing episodes based on novel sensory combinations. Despite its great allure, the subjective nature of this covert experience and the lack of clear observable behavioral markers make its experimental investigation notoriously difficult (Hubbard, 2018). Thus, despite progress in showing similar global neural responses to perceived and imagined events, we do not really understand how the actual contents of internal sensory experiences are represented in the brain, particularly in the case of auditory imagery, which has been less studied than visual imagery. Are the neural codes elicited by perception and imagery of the same auditory event similar? Do they represent the temporally unfolding auditory content in our internal thought?

At a phenomenological level, many sensory and cognitive aspects of perceived sounds are reported to also be experienced in imagery, and imagery of music is no exception ((Lucas et al., 2010), see (Hubbard, 2010) for a review). Yet the neural realization of these similarities is not well specified. We know that the general topography of brain areas recruited when people imagine and perceive musical sounds is similar. Bilateral auditory cortices are consistently recruited in both imagery and perception of sounds (Zatorre and Halpern, 1993; Griffiths, 2000; Halpern, 2001; Halpern et al., 2004; Kraemer et al., 2005; Herholz et al., 2012; Ding et al., 2019), as are frontal and parietal cortical structures such as the supplementary motor area (Zatorre et al., 1996; Halpern and Zatorre, 1999; Hickok et al., 2003; Foster and Zatorre, 2009; Herholz et al., 2012; Foster et al., 2013; Ding et al., 2019). Thus, prior studies have clearly established that individuals are able to voluntarily reinstate musical content in their “mind’s ear”, and that there is a shared neural substrate for perception and imagery. However, it is still a mystery whether these neural substrates represent the *content* of experience, regenerated during imagery and evolving with the flow of the conscious mind (James, 1890), or rather only share a global involvement in both imagery and perception.

The neural representation of internal reinstatement of complex sounds has been most commonly studied by looking for spatially overlapping neural responses while listening versus imagining sounds. Yet a brain area could respond reliably to both internal and external sounds without the same operations necessarily being carried out in each case, and with the same temporal dynamics (Dinstein et al., 2007; Ben-Yakov et al., 2012; Regev et al., 2019). Such findings are therefore limited in terms of differentiating between representations of specific imagined content reinstated from memory (e.g., one sound vs. another). More recently, studies have investigated the different information that might be represented within a given region of interest during auditory imagery, using decoding methods. Most of these efforts focused on spoken language, making great progress in localization of the content-specific representation of imagined linguistic constructs, ranging from single vowels to full sentences (e.g., (Pei et al., 2011; Ikeda et al., 2014; Martin et al., 2014; Rampinini et al., 2017; Cervantes Constantino and Simon, 2018; Musch et al., 2019)). Nevertheless, the dominant semantic context and linguistic knowledge relevant for these paradigms can strongly influence the recruited sensory circuits that are involved in auditory imagery (Kraemer et al., 2005). It could be argued that when using internal speech, the identified neural representations, especially outside early auditory cortices, might in fact capture more abstract or symbolic processes unique to language, rather than reinstatement of auditory experience per se.

Other studies used imagined environmental sounds to identify spatially distributed neural representations, mostly in associative auditory cortices ((Meyer et al., 2010; Vetter et al., 2014; Linke and Cusack, 2015) but see (de Borst et al., 2016) for findings in somatomotor areas). While the ability to decode which environmental sound a person is imagining from fine-grained spatial patterns of cortical activity does not involve linguistic mechanisms present during internal speech, such findings do not address two critical issues. First, it is not clear if the decoded neural representations are the same during imagery as during listening to sounds, since direct evidence for reinstatement (during imagery) of neural patterns characteristic of auditory perception has been missing. Indeed, in contrast to findings in the visual system (Thirion et al., 2006; Albers et al., 2013), attempts to identify any reinstated neural patterns in auditory brain regions during imagery have either failed, or have observed them only in the somatomotor cortex (Vetter et al., 2014; Gu et al., 2019). Second, environmental sounds are mostly brief by nature and are thus missing the temporal dynamics that occur over an extended period of time. Given that humans rely heavily on complex auditory sequences for communication via speech and music, neural studies of environmental sounds cannot fully capture the temporal complexity and structural dependencies of auditory content crucial to human cognition. It therefore remains important to test whether the same neural populations share the representation of sound sequences across perception and imagery, and whether the temporal pattern of activity in these populations is associated with the specific non-linguistic content of complex sensory sequences.

Similarities in the representation of sounds across imagery and perception can be studied not only locally, but also by comparing interactions across neural populations. One of the most prominent neural attributes of complex sound perception is the interaction between the auditory and motor networks (Zatorre et al., 2007; Hickok et al., 2011; Morillon et al., 2015). The effect of movement on auditory perception and its related neural activity has been demonstrated mostly in the context of the modulation of self-produced sounds, such as speech or music, by associated movements such as articulation or musical performance (Zatorre et al., 2007; Rauschecker and Scott, 2009; Hickok et al., 2011; Repp and Su, 2013; Rauschecker, 2018; Reznik and Mukamel, 2019). Recent studies have shown that rhythmic tapping to ongoing external auditory stimuli can also modulate their perception by facilitating selection of relevant auditory events and refining the representation of pitch (Morillon et al., 2014; Nozaradan et al., 2016; Morillon and Baillet, 2017). These results suggest that even when movements are not the source of the perceived sounds, their co-occurrence is sufficient to induce modulation in the auditory response. Interestingly, there is evidence showing that rhythmic movement also affects the mnemonic representation of sounds. For example, tapping along during an initial exposure to music has been shown to increase its memorability and chances of becoming an earworm (Mikumo, 1994; McCullough Campbell and Margulis, 2015). These results, together with the consistent evidence for the neural coactivation of motor regions during auditory imagery (see (Lima et al., 2016) for a review) raise the possibility that the strong auditory-motor relationship might allow movement to modulate the internal representation of sounds.

In the present work, we compared the unique temporal neural response profile of imagined and heard musical pieces. This approach allowed us to probe the specific contents of imagined auditory experiences that unfold over time, and also to explore the influence of rhythmic motion on the neural representation of auditory imagery. Prior to scanning, participants memorized six different ~1 minute-long instrumental pieces, and the accuracy of their musical imagery was verified behaviorally. Next, brain activity was measured with functional magnetic resonance imaging as participants either listened to each of the memorized melodies, or imagined each melody under two conditions: with and without simultaneous rhythmic tapping to a visual metronome. In this manner, the timing of the auditory stimuli and imagery were precisely matched across conditions and participants, allowing us to measure temporal alignment of brain responses.

We used inter-subject correlation (ISC) analyses, which provides a tool for extracting neural activity that is locked in time to continuous natural stimuli (Hasson et al., 2004; Hasson et al., 2009). By measuring the reliability of neural response time courses across brains, we can detect neural activity that is specifically induced by temporally extended experiences shared by participants. For example, previous studies using this method have shown that listening to the same music (Alluri et al., 2012; Abrams et al., 2013; Farbood et al., 2015) or vocalizing the same learned story (Silbert et al., 2014) can induce strong between-participant similarity in the time courses of brain activity in many regions. We extend this approach to investigate neural commonalities between people during purely *internally driven* mental processes, covertly generated with no external manifestation. ISC analyses also allow us to measure ongoing components that are shared across different experimental conditions, akin to the way the neural response to a narrative has been compared across different sensory forms of presentation (Honey et al., 2012; Regev et al., 2013; Nguyen et al., 2019). Here, we used this method to ask whether the shared content of inner conscious experience elicits a similar neural code across people, and whether this code is common to the external perceptual experience as well. Such a finding could demonstrate how the subjective experience of our inner and outer landscape is largely based on a common set of principles.

Thus, we first tested the prediction that imagery of auditory content can evoke similar response patterns across people, despite the lack of external input. Next, by comparing temporal response patterns across perception and imagery, we investigated whether responses during perception were reinstated during imagery in a melody-specific manner, allowing us to test the hypothesis that the specific contents of the conscious experience could be mapped. Finally, we studied the effect of rhythmic tapping on the reinstatement of melody-specific activity, to test the idea that top-down motor signals enhance internal auditory representations.

## Results

Twenty-five participants memorized six instrumental musical pieces, with two subgroups of three pieces matched for their beat timing, creating two tempo sub-groups (73 or 107 beats per minute; BPM) (see Methods, *Stimuli*). After an independent learning phase, participants were screened to ensure they had indeed memorized all the melodies. Participants were tested for their ability to accurately imagine the music, and then underwent a functional MRI session during which they performed the following conditions for each melody: (1) passively listening to the melody (“perception”); (2) silently imagining the music to the rhythm of a visual metronome – a bouncing ball at the center of the screen – while keeping motionless (“imagery w/o tap”); and (3) silently imagining the music to the rhythm of the metronome while tapping their finger to the beat of the music (“imagery w/ tap”; see Supplementary Fig. 1). In addition, participants took part in two control conditions in which they watched the same metronome as in the imagery conditions while tapping to its beat (“control w/ tap”) or keeping still (“control w/o tap”), but were not asked to imagine the music (see Methods, *Experimental design*).

### Behavioral assessment of imagery accuracy

Prior to the scans, participants completed two tests aimed at assessing the accuracy of their imagery of each of the learned melodies. The “Beat” test and the “Pitch” test were designed to capture temporal and pitch fidelity of the internally recalled melody, respectively (Herholz et al., 2008; Weir et al., 2015). Each test was performed under two movement instructions: during imagery, participants were either asked to be still or tap their finger to the rhythm of the metronome (see Methods, *Experimental design*).

In both the Beat and Pitch tests, average success rate across the six melodies was higher than chance level (Beat: M = 72.3%, SD = 11.6%; *t*_(24)_ = 9.62, *p* ≪ 0.0001, *d* = 3.93; Pitch: M = 96.7%, SD = 2.4%; *t*_(23)_ = 94.97, *p* ≪ 0.0001, *d* = 39.6), demonstrating the participants’ ability to accurately internally recall the learned melodies (Supp. Fig. 2).

Next, we explored whether the accuracy of imagery was affected by rhythmic tapping. Average success rate was similar whether participants tapped or not during imagery in both the Beat test (tap: M = 72%, SD = 11.1%, no tap: M = 72.7%, SD = 15%; *t*_(24)_ = −0.27, *p* = 0.79, *d* = −0.11) and the Pitch test (tap: M = 96.8%, SD = 2.1%, no tap: M = 96.6%, SD = 3.4%; *t*_(23)_ = 0.36, *p* = 0.72, *d* = 0.15). With the confirmation of participants’ ability to accurately imagine the music, we examined whether there was neural reinstatement during imagery.

### Similar temporal patterns between participants during imagery

We began by establishing to what extent, and where in the brain, imagining the melodies elicited synchronized neural response across people. We identified the brain areas that responded reliably across participants who imagined the same melody (without tapping) by calculating the temporal inter-subject correlation (ISC; (Hasson et al., 2004)) within each of 400 cortical parcels (Schaefer et al., 2018) across brains (Fig. 1A). The response reliability map of imagery averaged across all six melodies is shown at r > 0.14 for visualization purposes, and a familywise error rate (FWER) was used to correct for multiple comparisons (Fig. 1B).

**Figure 1.**
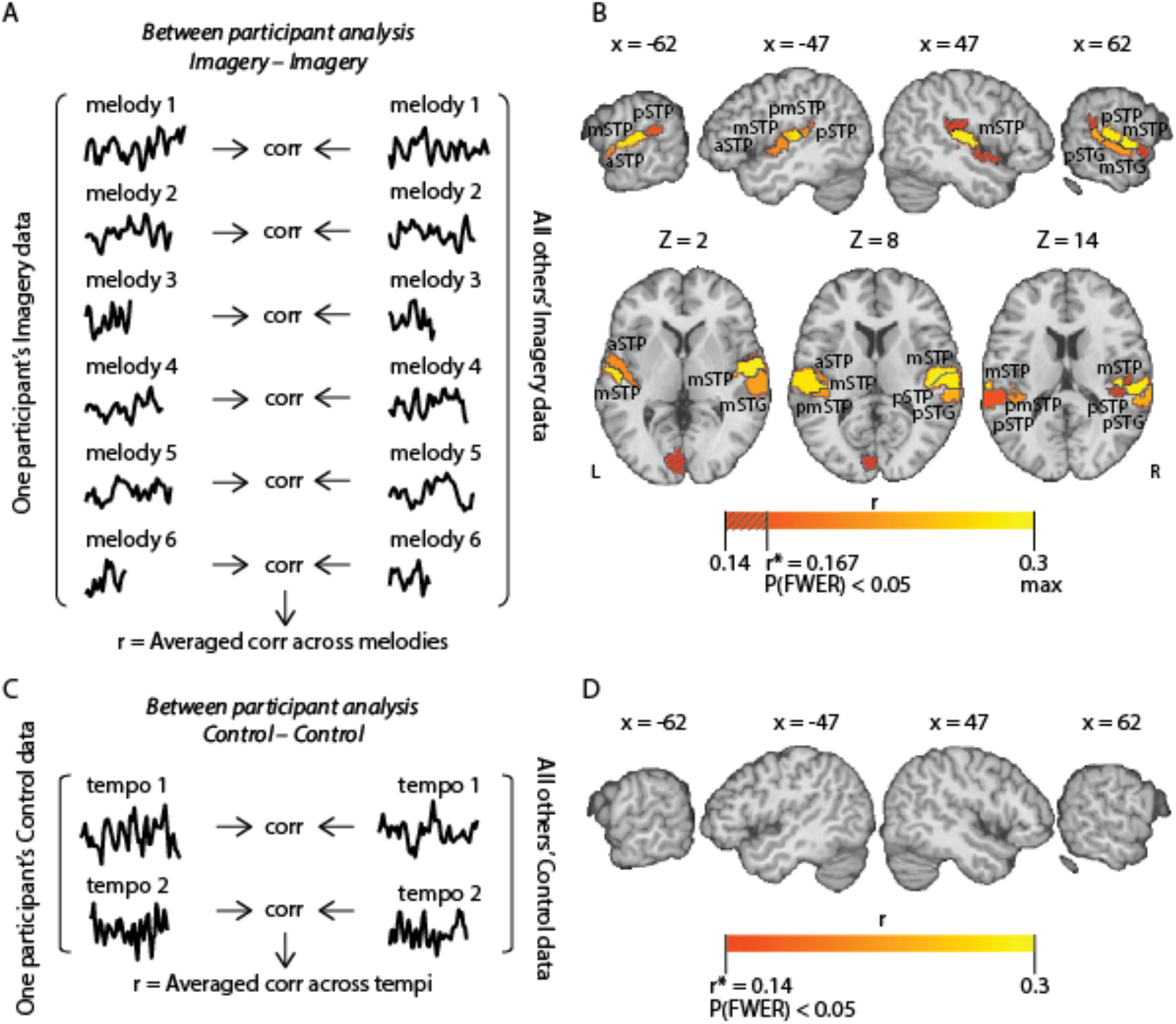
Temporal pattern similarity in the imagery and the control conditions (w/o tap). **A,** Schematic for between-participant analysis for the imagery condition. BOLD time courses were correlated between participants (each participant vs. the average of all others) for each of the six melodies in each parcel, to produce an averaged ISC during the imagery task. **B,** Cortical parcels where the highest temporal pattern similarity was observed across participants during imagery of the same melodies (*p*_(FWER)_ < 0.05; threshold r = 0.14 for visualization purposes). Regions in the lateral STG include middle and posterior parcels (mSTG, pSTG); Regions in the STP include middle, posterior, anterior and posterior-medial parcels (mSTP, pSTP, aSTP, pmSTP). **C,** Schematic for between-participant analysis for the control condition, same as A, except that correlations were computed for each of the two types of task tempi. **D,** No cortical parcels in the temporal cortex show significantly similar temporal patterns between participants (*p*_(FWER)_ < 0.05).

We found similar temporal responses across participants during imagery of each of the melodies in early and associative auditory cortices (Fig. 1B). These areas included parcels at the middle of the superior temporal plane (mSTP) bilaterally, which largely overlap with loci of early auditory processing in Heschl’s gyri (Supplementary Table 1), as well as in more anterior and posterior auditory areas in the STP, and in lateral parts of the right superior temporal gyrus (STG). Most of these sensory regions also showed similar temporal response across participants during the perception of the melodies (Supp. Fig. 3A & B).

To exclude the possibility that the observed similarities in the temporal response patterns were driven by the visual metronome (which was present in the imagery condition), rather than the imagery task itself, we performed the same inter-subject correlation analysis in the control condition (Fig. 1C), in which participants were exposed to the same sensory input as in the imagery condition (a bouncing ball in one of two tempi) but were not asked to imagine the melodies. Although some involuntary imagery might have taken place during the control condition, this analysis nonetheless revealed no significant correlations in the auditory cortices (Fig. 1D), thus confirming that the similar temporal patterns across people are driven by the mental imagery task, rather than the visual input.

### Similar temporal pattern across perception and imagery

The preceding inter-subject correlation in responses during imagery could be attributed to any of several different components of the imagery task. It might have been evoked by a common reinstatement of the previously perceived experience of music, but could also be due to nonspecific shared processes that are idiosyncratic to imagery, which are not manifested during music perception (e.g., certain control mechanisms or systematic modified mnemonic representation of the melodies). To investigate this question, we next looked for areas that specifically took part in the reinstatement of the perceptual experience of the melodies during imagery.

Although spatial overlap between areas that are activated during both auditory imagery and perception suggests some level of shared neural processing, a stronger form of shared neural representation is indicated when a region responds with the same temporal response profile to imagined and perceived forms of the same specific musical content. To test for direct correspondence between perception and imagery, we correlated the response time courses within each parcel when a given melody was perceived, to the response time courses when it was imagined. This analysis was performed on a between-participant basis by comparing each participant’s response during imagery to that of all others during perception, which, when averaged, served as a group template of the typical response pattern for the perceptual experience (Honey et al., 2012; Chen et al., 2017; Nguyen et al., 2019; Regev et al., 2019) (Fig. 2A). This analysis revealed that in the bilateral early auditory areas (mSTP) and other higher auditory areas in the right STP and STG (Fig. 2B), temporal response patterns obtained during imagery of a given melody were similar to the patterns in other individuals during perception of that same melody. Thus, the results indicate a shared neural response across perception and imagery, which is locked to the temporal dynamic of the melodies and consistent across individuals.

**Figure 2.**
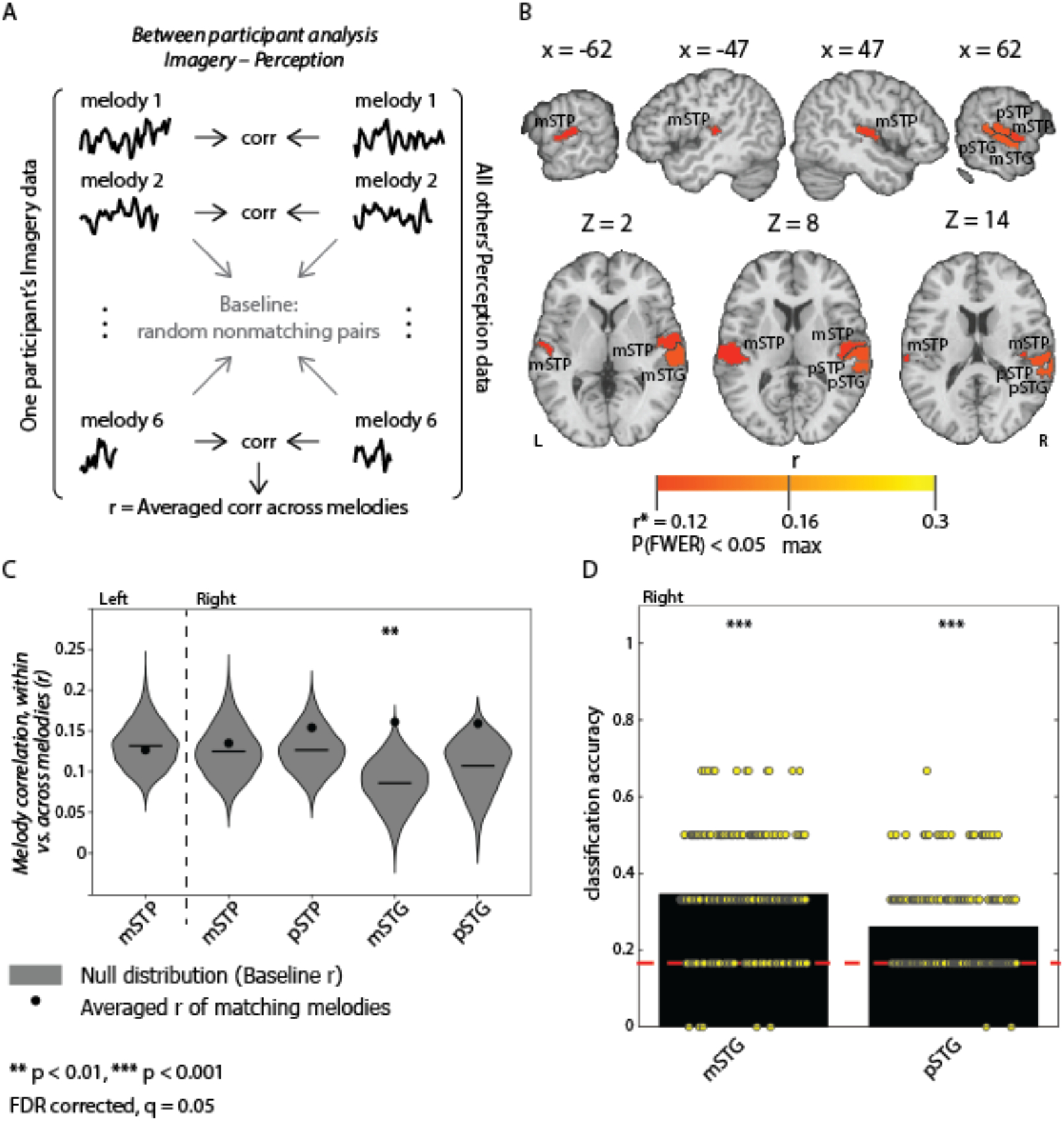
Temporal pattern similarity between imagery (w/o tap) and perception. **A,** Schematic for between-participant similarity analysis across imagery and perception. The correlations were computed between every matching (black) and non-matching (grey) pair of imagery/perception melodies across participants (non-matching comparisons are used for the analysis described in C). **B,** Cortical parcels where the highest temporal pattern similarity was observed across participants between imagery and perception of matching melodies (*p*_(FWER)_ < 0.05). **C,** Discriminability of temporal patterns in parcels that showed similarity across imagery and perception. Reinstated temporal patterns were discriminable from each other when the similarity between matching melodies was greater than between non-matching melodies. Black circles show average correlation of matching melodies. Statistical significance was determined by generating a null distribution of random pairs of non-matching melodies (grey violin baseline; FDR-corrected *q* = 0.05). **D**, Classification accuracy of melodies across brains between imagery and perception in the right mSTG and pSTG. Participants were randomly assigned to one of two groups (N = 12 and N = 13), and an average time course calculated from the imagery condition in one group and from the perception condition in the other for all six melodies. Accuracy was calculated as the proportion of melodies correctly identified out of six. The entire procedure was repeated using 200 random combinations of the two group sizes (yellow marks), and an overall average was calculated for each parcel (right mSTG 35%, pSTG 26%, black bars; chance level 16.7%, red line; FDR-corrected *q* = 0.05).

A comparison of the response patterns in perception to the responses in the control condition revealed no significant similarities in the auditory cortices (Supp. Fig. 4A & B), suggesting that the mere presence of the visual rhythm of the metronome did not account for the reinstatement of the neural patterns during imagery.

Does high similarity in temporal responses between imagery and perception of melodies reflect an accurate internal representation of the melody? To answer this question, we compared each individual’s magnitude of inter-subject imagery-to-perception similarity, averaged across songs, to their general score in the behavioral Beat test of imagery accuracy. The correlation between these two scores revealed the strongest associations to be in the right middle and posterior STG (*r* = 0.4, *p* = 0.047 and *r* = 0.43, *p* = 0.037, uncorrected for multiple comparisons; Supp. Fig. 5), out of the all parcels where the similarity of response between imagery and perception was significant. This might indicate that the reinstatement of neural responses during imagery in these regions in the right STG can predict the accuracy of the internal musical experience.

### Neural reinstatement of melody-specific temporal patterns

The consistency in temporal response pattern across perception and imagery could potentially represent general cognitive trends common in both tasks, regardless of the identity of the melody (e.g., rapid buildup of attention). At the same time, the similarities between perception and imagery could also represent the reinstatement of the specific musical content, and be driven by temporal response patterns that are unique to the experience of each melody. To test whether the reinstated responses are discriminable from each other, we used a permutation analysis (Kriegeskorte et al., 2008; Baldassano et al., 2018) that compares neural pattern similarity between matching melodies against the nonmatching melodies, across perception and imagery (see Methods, *Discriminability analysis*). This analysis reveals regions containing reinstated melody-specific patterns, as statistical significance is only reached if matching melodies (same melody in perception and imagery) can be differentiated from nonmatching melodies (Fig. 2A & C, grey baseline). The discriminability analysis was performed in each of the five parcels in the STP and STG that showed a significantly similar response pattern across perception and imagery. The results of this analysis showed a within-vs-between melody difference in the right mSTG (*p* < 0.005, FDR corrected *q* = 0.05), and a smaller difference in the right pSTG (did not survive correction; Fig. 2C).

To further explore *how* discriminable the reinstated neural patterns of melodies were from each other, we performed a classification analysis (Chen et al., 2017) for the parcels that showed melody-specific response patterns (i.e., right mSTG and pSTG). Participants were randomly assigned to one of two groups (*N* = 12 and *N* = 13), and an average time course for each melody was calculated from the perception condition in one group and from the imagery condition in the other group. Classification accuracy across groups was calculated as the proportion of melodies correctly identified out of six. The procedure was repeated using 200 random combinations of groups and averaged. Statistical significance was assessed using a permutation analysis in which the melody labels were randomized before calculating the accuracy (see Methods, *Classification accuracy*). The classification accuracy was found to be significant in both the right mSTG and pSTG, with 35% and 26% correctly labeled melodies respectively (Fig. 2D; chance level 16.7%, *p* < 0.001; FDR corrected *q* = 0.05).

Since the two different tempi of the melodies may have contributed to the discriminability of the neural patterns between melodies, we repeated the classification analysis within each tempo sub-group of 3 melodies. The classification accuracy was again found to be significant within each of the two tempo sub-groups at the right mSTG (BPM 73: 44%, BPM 107: 45%, *p* < 0.001; chance level: 33%), but only in one sub-group at the right pSTG (BPM 73: 41%, *p* < 0.001; BPM 107: 34%, *p* = 0.32; Supp. Fig. 6A & B). Overall, this suggests that the right mSTG, and perhaps less so the right pSTG, reinstate melody-specific temporal neural patterns during imagery. These specific patterns are not unique to the internal experience of individuals but are shared across all participants who imagine the same musical content.

### The effect of rhythmic tapping on reinstatement of melody-specific temporal response patterns

Do neural representations of melodies change when imagery is accompanied by rhythmic movement? To address this question, we examined the effect of rhythmic tapping to the beat of imagined melody on the reinstatement of its unique response pattern, and on the propagation of the imagined content across functional networks.

For this purpose we began by repeating the ISC analysis, which compared the response time courses when a melody was perceived, to the response when it was imagined, but with accompanying rhythmic tapping movements (Fig. 3A). The analysis revealed similar temporal response patterns between perception and imagery, which extended to further auditory areas, beyond parcels identified in the previous comparison to motionless imagery. Specifically, more parcels in the STP bilaterally and in the left lateral STG showed significant response similarities across perception and imagery of melodies due to the added motor component to the imagery task (Fig. 3B).

**Figure 3.**
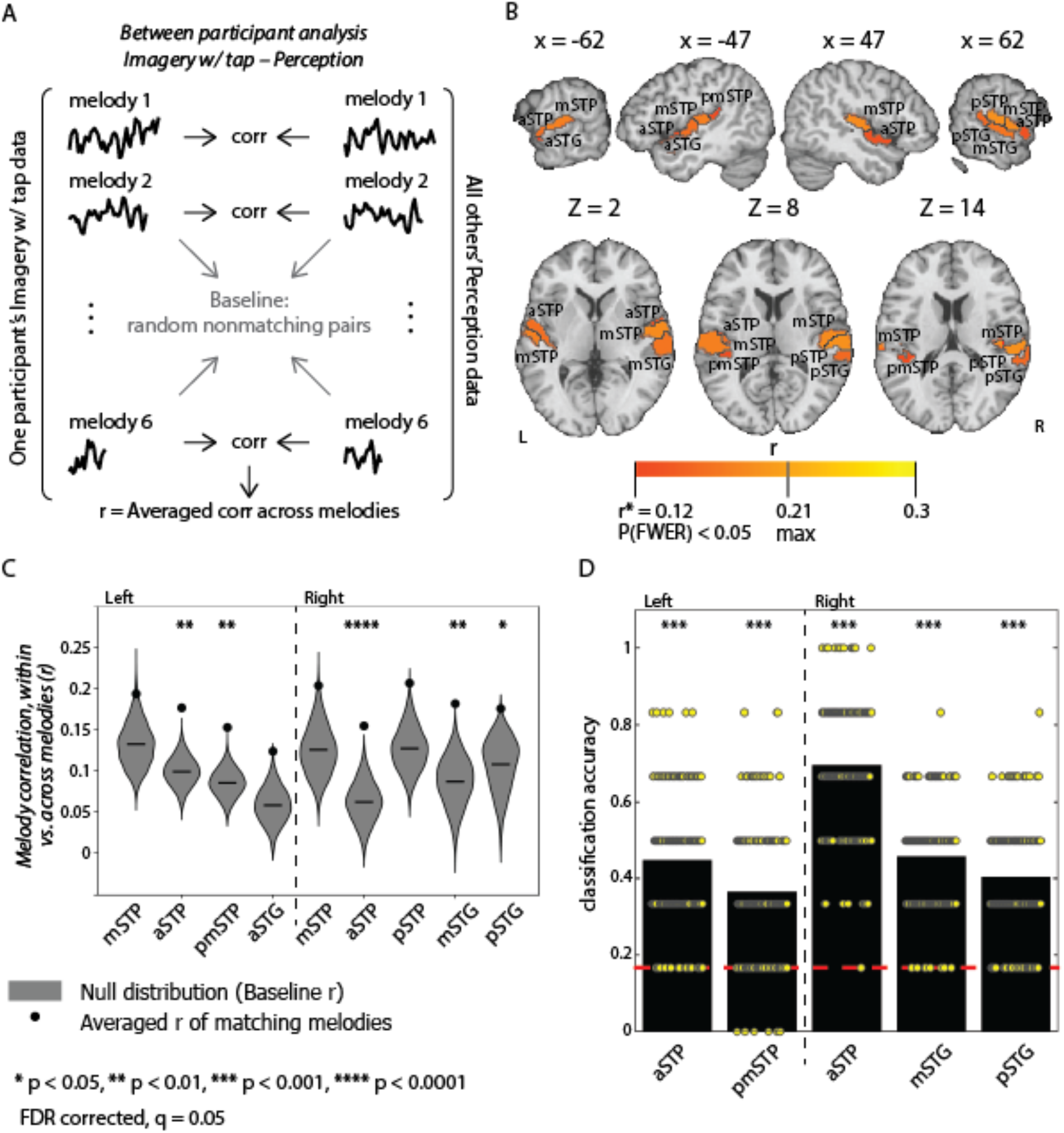
Temporal pattern similarity between imagery accompanied by tapping and perception. **A,** Schematic for between-participant similarity analysis across imagery w/ tap and perception. Same as Fig. 2A, except that correlations were computed using the imagery condition that included tapping. **B,** Cortical parcels where the highest temporal pattern similarity was observed across participants between imagery w/ tap and perception of matching melodies (p_(FWER)_ < 0.05). **C,** Discriminability of temporal patterns in parcels that showed similarity across imagery w/ tap and perception (FDR-corrected = 0.05) (see Fig. 2C). **D,** Classification accuracy of melodies across brains between imagery w/ tap and perception (right mSTG 46%, pSTG 40%, aSTP 70%, left aSTP 45%, pmSTP 36%, black bars; chance level 16.7%, red line; FDR-corrected q = 0.05) (see Fig. 2D).

The introduced synchronized movements also enhanced the reinstatement of melody-specific response patterns in the auditory cortex. The discriminability analysis was performed using the movement-accompanied imagery data in the nine parcels in the STP and STG that showed significantly similar responses across perception and imagery. We identified significant melody discriminability in regions that also exhibited such response patterns during motionless imagery (right mSTG *p* < 0.005, right pSTG *p* < 0.05; Fig. 3C), as well as an increase in classification accuracy (right mSTG: 46%, *p* < 0.001; *t*_(398)_ = 7.77, *p* ≪ 0.0001, *d* = 0.78; right pSTG: 40%, *p* < 0.001; *t*_(398)_ = 10.33, *p* ≪ 0.0001, *d* = 1.04; Fig. 3D). Furthermore, the discriminability extended to further regions in the STP bilaterally, including the aSTP parcels (left *p* < 0.005, right *p* < 0.0001) and left pmSTP (*p* < 0.001), which also showed significant classification accuracy (left aSTP 45%, right aSTP 70%, and left pmSTP 36%, *p* < 0.001). All these regions, except the right pSTG, showed significant classification accuracy as well, when calculated within each of the different tempo sub-groups (Supp. Fig. 6C & D). This suggests that temporal patterns of brain activity in the associative auditory cortex (especially parts of the STP) were modified by the incorporation of synchronized tapping with imagery in a consistent manner across individuals, in becoming more similar to responses during perception, and more distinguishable across melodies. Thus, tapping did not just improve the general beat representation of the melodies but also strengthened the unique neural representation of melodies with the same tempo.

### The effect of rhythmic tapping on propagation of musical content across cortical areas during imagery

What functional connections in the brain might support the observed spread of melody-specific information during rhythmic movement? To map how melody-locked neural responses are shared across pairs of brain areas, we used inter-subject functional correlation (ISFC) analysis (Simony et al., 2016). In this analysis, the inter-regional comparison of temporal responses was performed on a between-participant basis, and across the perception and imagery conditions. By measuring the correlation across brains, instead of within a brain, the intrinsic neural correlations and non-neural confounds can be filtered out, thus increasing the ability to detect coupling induced by shared experiences across participants. In addition, by comparing the responses across conditions, we can identify couplings driven by the commonalities across perception and imagery. This analysis was performed for each melody between the perception and each of the two imagery conditions (w/ or w/o tapping) across all 400 parcels, which were organized in resting-state functional networks ((Yao et al., 2011; Schaefer et al., 2018); Supp. Fig. 7A & B).

To describe the degree of inter-regional coupling enhancement by tapping, we calculated the difference of the mean correlation during motion-accompanied imagery from the mean correlation during motionless imagery. The more positive the resulting value, the more reliable the inter-regional responses while tapping, whereas negative values indicate that the responses are more reliable without tapping. Significant enhancement of coupling was determined using a (paired two-tailed) permutation analysis that randomly assigned the correlations from the two imagery conditions (with or without tapping) to two new sham groups, generating a null distribution from the subtraction between the sham groups. We corrected for multiple comparisons across all 400 parcels by controlling the FDR (*q* = 0.01).

Melody-locked inter-regional responses were most strongly enhanced between parcels of the ventral somatomotor (VSM) network when tapping accompanied imagery (M = 0.048, SD = 0.017; Fig. 4A, pink). This functional network encompasses the STP (Fig. 4B and C), including the parcels in which melody-specific reinstatement was extended to when imagery was accompanied by tapping. The response enhancement in this network was not limited to these auditory parcels but also extended between the STP and insular and ventral somatomotor areas.

**Figure 4.**
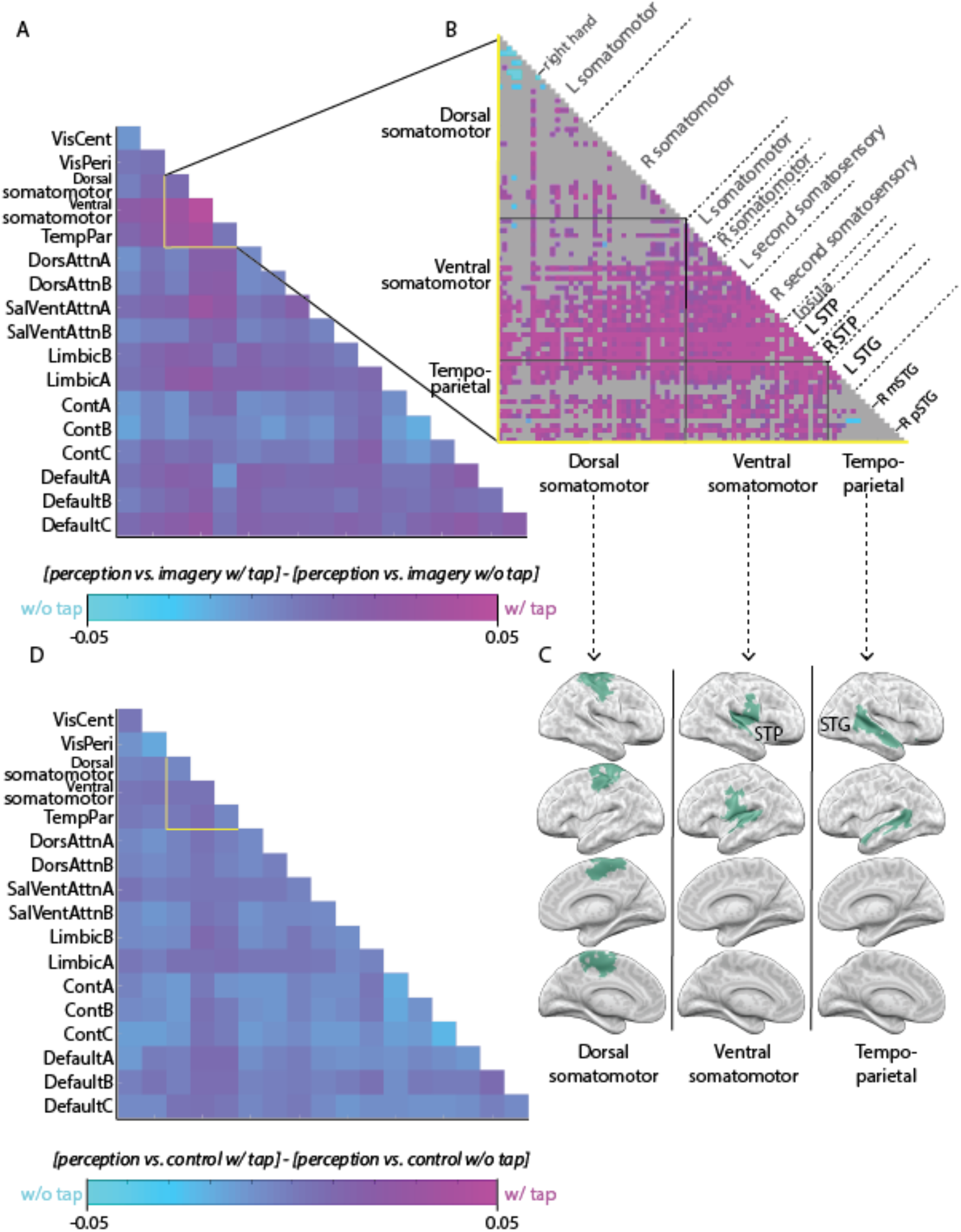
Rhythmic tapping during music imagery enhanced inter-regional melody-locked responses across functional networks. Degree of enhancement was assessed as the difference in level of melodies’ reinstatement when imagery was accompanied with tapping and was motionless. **A,** Inter-regional coupling in reinstatement of music was enhanced across several functional networks. **B,** The strongest enhancements were observed between parcels in the ventral somatomotor network, and between this network and parcels in the tempo-parietal and dorsal somatomotor networks (FDR-corrected q = 0.01). **C,** The ventral somatomotor network includes the parcels in the STP, as well as ventral somatomotor areas. The tempo-parietal network includes parcels in the lateral STG, and the dorsal somatomotor includes parcels which corresponds to hands topography (Yao et al. 2011). **D,** The enhancement of inter-regional coupling by tapping was low when assessed as the difference between the control conditions w/ and w/o tap.

Furthermore, tapping enhanced melody-locked inter-regional responses between the VSM network – especially the STP – and other functional networks. We observed the strongest increase in inter-network coupling with the temporo-parietal network (M = 0.036, SD = 0.016; Fig. 4C), including parcels in the right STG that showed melody-specific reinstatement during imagery (Fig. 2C and 3C), and with the dorsal somatomotor (DSM) networks (M = 0.033, SD = 0.021; Fig. 4B), which was previously found to accommodate the hand and foot regions ((Yao et al., 2011); Fig. 4C). Other strong coupling enhancements were observed between the VSM network, especially the STP, and cognitive networks, such as one of the ventral attention networks (M = 0.03, SD = 0.019), as well as with the peripheral visual cortices (M = 0.025, SD = 0.015) (Fig. 4A).

To further test whether the more reliable inter-regional responses while tapping during imagery were indeed driven by the musical experience common across perception and imagery, we repeated the inter-regional analysis using the data from the control conditions instead of the imagery conditions (Supp. Fig. 7C & D). The subtraction between the correlations during motion-accompanied and motionless control showed a relatively weak enhancement in inter-regional coupling during tapping across all functional networks (e.g., in VSM M = 0.004; Fig. 4D). These results demonstrate that the enhancement in coupling during tapping-accompanied imagery is not likely to be an artifact simply due to the inclusion of motion in the task, but rather represents an increase in the propagation of information related to the musical imagery task itself.

## Discussion

We found that the specific contents of consciousness during auditory imagery can align the neural responses across individuals in both early and associative auditory cortices (Fig. 1). In a subset of these regions, we found that the temporal response patterns recorded during the perception of music were reactivated during imagery in a melody-specific manner, indicating that the response is specific to the imagined content. This reinstated and shared brain activity was observed in the middle of the right superior temporal gyrus, in the absence of any auditory sensory cues (Fig. 2). Furthermore, when music imagery was accompanied by synchronized rhythmic tapping, the distinctive responses to the melodies were reinstated in additional associative auditory areas of the superior temporal plane (Fig. 3), thus increasing the resemblance to the neural representation of the perceived stimulus. This extended reinstatement was also accompanied by an increase in the spread of the melody-locked response between the STP and varied functional networks, most prominently with somatomotor areas (Fig. 4), all of which supports an important role for motor-to-auditory influences in enhancing the fidelity of internal auditory representations.

### Neural representation of specific imagined musical content

Prior studies reported spatial overlap between areas that respond to perceived and imagined music (e.g., (Halpern and Zatorre, 1999; Hickok et al., 2003; Kraemer et al., 2005; Meyer et al., 2007; Herholz et al., 2012)). However, such overlap does not reveal whether the region contains information about the specific ongoing auditory content being retrieved and experienced. The present study goes beyond previous findings by demonstrating that brain activity time courses when listening to real-life complex music were reinstated during imagery of the music in the STP and STG (Fig. 2B), and that these reinstated melody patterns were discriminable from one another in the right STG (Fig. 2C & D). Moreover, our methodology revealed the extent to which the subjective internal experience of hearing music in the “mind’s ear” has shared neural representations across people, even though each person learned the melodies as they chose, and had different levels of musical expertise.

The similarity in temporal responses across perception and imagery was observed in early auditory areas bilaterally (Fig. 2B). Previous studies have found a spatial overlap in neural activity in secondary and associative areas, but evidence for shared activity is scarcer for primary areas (e.g. (Griffiths, 2000; Halpern et al., 2004; Bunzeck et al., 2005; Herholz et al., 2012), but see (Yoo et al., 2001; Kraemer et al., 2005; Oh et al., 2013)). While between-participant synchronization in the early auditory cortex has been reported during perception (Lerner et al., 2011; Honey et al., 2012; Regev et al., 2013; Farbood et al., 2015), this is the first study to capture temporal alignment in this sensory area during imagery, a purely internal process without any externally driven acoustic component. Although we observed similarities in parcels that overlap with Heschl’s gyrus, the whole-brain parcellation used here (based on resting-state functional connectivity; (Schaefer et al., 2018)) does not allow an accurate dissociation between primary and secondary regions. Therefore, it is possible that the recorded similarities in the early auditory cortices are driven by activation in the secondary, rather than primary auditory regions.

Despite the similarity between perception and imagery in the early auditory areas, we did not identify a reinstatement of melody-specific responses during imagery in these areas, but rather in lateral areas of the STG (Fig. 2C). This result is consistent with previous neuroimaging studies that attempted to decode environmental sounds across perception and imagery using multi-voxel pattern analysis, but did not find distinctive representation in early auditory areas (Meyer et al., 2010; Vetter et al., 2014; de Borst et al., 2016; Gu et al., 2019). Our results suggest that the temporal response similarity observed in the early auditory region across perception and imagery might represent general operations shared by both tasks, with temporal dynamics that are not specific for each melody (but might still influence the auditory experience).

Previous studies have shown that during processing of real-life auditory stimuli, such as music and speech, activity time courses in associative auditory regions are synchronized across individuals and locked to mid-level structures in the stimulus (e.g. sentences and musical phrases; (Lerner et al., 2011; Farbood et al., 2015)). The current study extends these findings by showing that synchronized neural responses during perception can give rise to shared neural responses during imagery in the right lateral associative auditory cortices, reinstated at will from memory (Fig. 2C & D). Although such neural reinstatement during auditory imagery was never captured before, this finding corresponds well with previous studies that revealed neural organization in high-level visual areas for recalled memories of specific movie scenes, that is similar to perception and sometimes common across participants (Buchsbaum et al., 2012; Chen et al., 2017; Zadbood et al., 2017). In agreement with our findings in the auditory modality, the key role of the right auditory cortex in imagery of music has been described before in temporal-lobe patients and noninvasive neuroimaging studies (Zatorre and Halpern, 1993; Halpern and Zatorre, 1999; Meyer et al., 2007; Ding et al., 2019), and is also in keeping with an extensive body of evidence supporting the idea that melodic information is better represented in auditory areas within the right vs. left hemisphere (Zatorre et al., 2002; Albouy et al., 2020).

In general, the responses observed here during either imagery or perception were limited to the temporal lobe. During imagery of music, neural activity has previously been described also in frontal regions and most frequently in the supplementary motor cortex (Hickok et al., 2003; Leaver et al., 2009; Herholz et al., 2012; Jacobsen et al., 2015). Although we did not observe any significant response in these areas, the lack of reliable dynamic responses across people does not imply there was no neural activity in these regions, only that this pattern of activity was not time-locked to the common imagined content. It is possible that additional regions were active during imagery and perception, but did not respond dynamically to the presented or retrieved information (for example, they could have maintained a constant level of activity over time, related to more general cognitive or sensory processes).

Unlike the current study, a reliable inter-subject temporal response during music perception was previously captured in structures in the frontal and parietal lobes (Alluri et al., 2012; Abrams et al., 2013; Farbood et al., 2015). However, these structures varied across experiments, and the response reliability there was weaker than described for the temporal cortex. This discrepancy with our results, which showed mainly the superior temporal cortex (Supp. Fig. 3B), could be a result of differences in stimulus and task demands. For instance, we used shorter musical segments (a single minute instead of longer melodies), which was essential for allowing their memorization, but cannot reveal areas that encode information integrated over longer periods of time (Farbood et al., 2015).

### The effect of rhythmic movement on the representation of imagined music

Temporal response patterns in the auditory cortex during musical imagery became more similar to the responses during perception when rhythmic tapping accompanied the imagery. Since the only shared experience across imagery and perception was the representation of music, and not the visual metronome or rhythmic movement (which were present only during imagery), the observed response similarities cannot directly represent any potential propagation of the rhythmic oscillations from visual or motor cortices into the auditory cortex. In addition, when the responses during perception were compared with a parallel control condition to imagery that included tapping to the metronome without an internal representation of music, no significant signal correlations were observed in the brain (Supp. Fig. 4C & D). Thus, although the spread of shared responses further into the STP could have been facilitated by a rhythmic modulation of the auditory cortex (e.g. through entrainment), this simple rhythmic component is not the source of the observed cross-condition similarities.

The strongest increase in neural similarity to perception and melody-specific reinstatement was observed in bilateral associative auditory areas in the STP, anteriorly and posteriorly to early auditory cortex (Fig. 3). These associative auditory areas in the temporal plane might differ from those in the lateral STG in their involvement in integration and recognition of relatively more fine-grained auditory patterns (Binder et al., 2000; Woods et al., 2009; Norman-Haignere et al., 2015; Hamilton et al., 2020), yet the distinct functional role of these regions in humans is still under debate. Thus, the movement might have enhanced various aspects of sensory processing that take part in both perception and imagery of music, such as spectrotemporal tuning of the neural receptive fields (Martin et al., 2018), or the sharpening of temporal selection of relevant auditory information (Nozaradan et al., 2016; Morillon and Baillet, 2017).

The increase in the similarity of the responses between perception and imagery and in the reinstatement of specific melodies could have been driven by modulation of different components of the temporal signal. For instance, the tapping manipulation might have had a broad effect on the memory of the melody as a whole, making the neural response during imagery more similar to perception in a uniform manner over time. Alternatively, the enhancement might have had a more oscillatory profile, with focal increases in similarity to perception around the co-occurring tapping events (Morillon and Baillet, 2017). Furthermore, the increase in similarity of the responses during imagery to those during the original sensory experience might have captured different enhancements in the internal representation of music. It could have been a result of more accurate neural representation of certain auditory features, instead of or in addition to participants’ possible improved ability to align their imagery to the metronome. If only the latter is true, inter-subject correlations might have increased without any rise in a population’s SNR of the perceptual representation. In this hypothetical case, our results would suggest that the added rhythmic movement had an ability to increase the temporal accuracy of retrieval of music, compared to a unimodal visual presentation of rhythm (as introduced by the visual metronome).

The strong anatomical and functional connection between the somatomotor and auditory cortices has been widely reported in variety of contexts, including both comprehension and production of complex auditory content (Hickok et al., 2003; Zatorre et al., 2007; Saur et al., 2008; Peretz et al., 2009; Rauschecker and Scott, 2009; Hickok et al., 2011; Rauschecker, 2011; Andoh et al., 2015). The strong interaction between these systems was specifically demonstrated in top-down somatomotor influences on the processing of perceived music, whether the movement generated the sounds (Zatorre et al., 2007; Repp and Su, 2013) or just accompanied them (Nozaradan et al., 2016; Morillon and Baillet, 2017). These influences were also demonstrated in the enhancing effect of articulatory movement on processing of the silently uttered speech (Okada et al., 2018; Zhang et al., 2020). To our knowledge, our results are the first to suggest that the motor system might have a broader modulatory role, not only during covert speech or when making sense of perceived sounds, but also when non-linguistic auditory information is represented entirely internally.

We found that the extended neural reinstatement of melodies in the auditory cortex due to tapping was accompanied by increased interactions between somatomotor and auditory cortices. Tapping during imagery induced the strongest increase in shared musical content between auditory parcels in the STP and somatomotor parcels around ventral parts of the central sulci (Fig. 4). These parcels create together a functional network typical for resting state (Yao et al., 2011), thereby providing a potential avenue for auditory-motor interactions (such as the current motor modulation) during various tasks. Although the influence of tapping was most prominent within this network, it also encompassed interactions between auditory and motor regions in other functional networks, including parcels in the lateral STG and the dorsal somatomotor cortex. Furthermore, tapping during imagery also led to the spread of content-specific coupling between the STP and other neural systems, such as parts of the ventral attention network and visual cortices. This suggest that the effect of rhythmic movement on internal representation of music might be a result of a complex interplay between motor, sensory, and high-order networks.

Whereas it is commonly hypothesized that motor commands lead to better prediction of sensory inputs and thus benefit individual’s adaptability to its environment (Wolpert et al., 1995; Schubotz, 2007; Schroeder et al., 2010; Cannon and Patel, 2020), the advantage of such a mechanism is not as clear during imagery, in which the sensory information had ostensibly already been established and can be retrieved at will. Nevertheless, the undeniable gap between stored knowledge and its transient representation in conscious experience could perhaps be somewhat mitigated by this mechanism. The modulatory capability of the motor system may be “repurposed” using deliberate motor strategies, in order to enhance the experience of covert auditory information, as has been shown before for overt speech and music (Kotz et al., 2009; Morillon et al., 2014; Schön and Tillmann, 2015; Morillon and Baillet, 2017). While in the current design tapping to the beat of the music was an imposed experimental manipulation, rhythmic movements often appear spontaneously when people imagine music, and silent articulation is common when seeking to establish internal speech during silent reading (McGuigan et al., 1964; Hardyck and Petrinovich, 1970). Thus, movement may have an important role in promoting the conscious reinstatement of complex auditory content.

Although this melody-specific reactivation was enhanced by rhythmic movement, we could not identify a corresponding behavioral improvement in imagery accuracy. Our measurement of imagery accuracy was modeled after an established measurement of internal pitch and beat representation (Herholz et al., 2008; Weir et al., 2015), but it was heavily modified for the use of longer and more complex musical segments, which could have influenced our ability to capture differences in imagery accuracy across conditions. As a result, we are unable to say how the tapping may have enhanced the internal experience of music by the participants, and the exact manner in which the modification of neural activity corresponds with that experience.

Overall, by following the neural responses to the specific content of imagery, this study was able to shed light on a largely impenetrable mental construct. The results reveal the existence of common neural activation for complex and continuous auditory content in regions where encoded information is largely sensory, bridging perceptual and imagined experiences. Over and above individual differences, we were able to trace the neural responses associated with familiar musical segments, and show the common ability to reactivate them at will when bringing to mind musical content. The fact that these observations were made as individuals were engaged with complex musical content testifies to the robustness and ecological validity of the phenomena. We also demonstrate that the similarity in the neural codes elicited by perception and imagery is not limited to isolated neural populations, but could also be observed in comparable intricate interactions between neural modalities, such as motor influence on the auditory system. Future work can explore what aspects of musical sequences (e.g., timbral, textural, or rhythmic) are most correlated with melody-specific fMRI time series during imagery (Alluri et al., 2012), and can study whether the objective and subjective mental reinstatement of sounds may be modified by the spread of neural reinstatement due to rhythmic movement.

## Acknowledgment

We thank Janice Chen and Benjamin Morillon for providing comments and suggestions, Vanessa Wong and the McConnel Brain Imaging Centre of the MNI for assistance in data collection, and to the Zatorre lab members for their support. This study was supported by a Canadian Institute for Advanced Research (CIFAR) Stimulus grant (to RZ, ADP and AMO), a Foundation Grant from the Canadian Institute of Health Research, (to RZ) and a postdoctoral fellowship from NSERC-Create in Complex Dynamics (to MR).

## Methods

### Subjects

Twenty-five participants were included in the final analysis (22 females, 3 males; age 18-32, mean age 21.9). Out of the initial 44 participants recruited, 17 were discarded for failing to memorize the melodies sufficiently, as assessed by the external-recall test (see *Learning phase*), and a further two were discarded from the analysis: one due to corrupted functional data and one due to head motion greater than 2mm.

Participants were selected without regard to their musical background, but were required to have learned at least 3 melodies and recalled them in the past by either vocalizing or playing an instrument. Their self-reported auditory imagery ability, as measured using the Bucknell Auditory Imagery Scale (BAIS), was not different than what has been previously reported in a larger sample of college students (current sample: M = 5.4 SD = 0.7; previous sample: *N* = 76, M = 5.1 SD = 0.9; unpaired two-tailed test: *t*_(99)_ = 1.52, *p* = 0.12; (Halpern, 2015)). All participants were right-handed, reported normal hearing and no absolute pitch, and did not report any neurological disorders. Procedures were approved by the Research Ethics Board of the Montreal Neurological Institute.

### MRI acquisition

Participants were scanned in a 3T full-body MRI scanner (Prisma, Siemens) with a 64-chanels head coil. Functional images were obtained with an interleaved multiband echo planer imaging (EPI) sequence (TE = 39ms; flip angle = 50°; multi-slice factor = 4; field of view [FOV] = 192 × 192 mm^2^; echo spacing = 0.55ms; 72 oblique axial slices), resulting in a voxel size of 2.0 mm isotropic and a TR of 1500ms. Anatomical images were acquired using a T1-weighted magnetization-prepared rapid-acquisition gradient echo (MPRAGE) sequence (TR = 2300ms; TE = 2.98ms; flip angle = 9°; 1.0 mm^3^ resolution; FOV = 256mm^2^).

### Stimuli

The auditory stimulus consisted of 6 distinct instrumental melodies composed by Joe Hisaishi for three different animated movies soundtracks: ‘Castle in the Sky’ (melodies [1] ‘Discouraged Pazu’ and [2] ‘Carrying You’), My Neighbor Totoro (melodies [3] ‘Evening Wind’ and [4] ‘Let’s Go to the Hospital’) and ‘Spirited Away’ (melodies [5] ‘One Summer’s Day’ and [6] ‘Reprise’). These melodies were written to evoke an emotional response in the audience, and exhibited full musical complexity, containing multiple instruments and melodic lines. At the same time, they also have a simple and catchy lead melodic line, making their memorization relatively easy. From each original track, we identified about a minute-long segment of the music ([1] 46 secs [2] 74 [3] 78 [4] 84 [5] 91 [6] 44), which contains clear and dynamic melodic line and a stable tempo.

The six melodies were split into two sub-groups, and the tempo of each three melodies was matched. To match the tempo, we first detected the beat locations for each original melody using the DBNBeatTracker from madmom library (Böck et al., 2016). Next, each melody was adjusted so that the average number of beats per minute matched the sub-group average (73 BPM for melodies 1, 3, & 5; 107 BPM for melodies 2, 4, & 6) and the beats were aligned (using the librosa library https://github.com/librosa/librosa/releases/tag/0.5.1). Thus, the melodies within each tempo sub-group shared an identical timing of beats. In addition, the waveforms of all melodies were matched for an averaged Root-Mean-Square value (implemented using librosa; the melodies can be found here: https://doi.org/10.5281/zenodo.3993675).

### Experimental design

#### Learning phase

Participants were asked to memorize the six melodies at home, during a period of 1-2 weeks, and spread their learning across at least two days. In addition to the melodies, they were provided with home-test videos that assisted in practicing accurate out loud recall of the melodies, in preparation for the memory test to come. These videos began with the first six seconds of the music, accompanied by a synchronized visual metronome in the shape of a white bouncing ball at the center of the black screen. After the music had stopped, the metronome carried on in silence, allowing participants to hum the rest of the melody to the beat. The music returned for the melody’s last 3 seconds, and thus provided participants with feedback whether their humming was correctly aligned with the original melody. The videos were accessed by participants through an online video player (Wistia.com), which enabled the experimenters to track the time participants spent listening to and recalling the melodies. As a condition for attempting the memory test and prove their knowledge, participants were asked to listen to each of the melodies at least 3 times, and use the home-test video at least 3 times successfully. Other than that, participants were not given any instructions regarding the strategy they should take for learning.

At the end of the learning period, a humming test was performed in the lab to assess participants’ ability to accurately recall the melodies. In this test, participants were shown the same video as used earlier during recall practice described above and were asked to hum the melodies out loud. A trained experimenter assessed the accuracy of their recall by evaluating the melodic contour and the temporal precision. Voicing quality, such as pitch accuracy or voice stability, was not considered, in an attempt to dissociate participant’s knowledge of the melodies from general singing abilities. To pass the test, participants had to fully recall each of the six melodies to the beat of the metronome, without skipping any beat. Successful participants were included in the experiment and continued to perform the internal memory test (see *Behavioral assessment of imagery*) and the fMRI scan.

#### Behavioral assessment of imagery

Before the scanning session, we assessed participants’ ability to accurately recall the melodies internally, without overt vocalization or lip movement. Each participant performed two memory tests (adopted from (Herholz et al., 2008; Weir et al., 2015)): (1) A “Beat” test, designed to capture accuracy of the recall of the melody in time, keeping the rhythm of the presented metronome without skipping a beat and (2) a “Pitch” test, designed to capture the ability to keep track of the original pitch of the internally recalled melody. Each test included three rounds in random order; each one assessed a different pair of melodies: [1]+[6], [2]+[3], or [4]+[5]. Each test was performed under two movement-instructions: during melody recall, participants were either asked to tap their right finger to the rhythm of a visual metronome or to stay completely still (considering any body movement, such of the finger, mouth, head or limbs).

##### Beat Test

Each trial in this test began with the first six seconds of the music, accompanied by a synchronized visual metronome in the shape of a white bouncing ball at the center of the black screen. After the music had stopped the metronome carried on in silence, allowing participants to imagine the rest of the melody to the beat. The music had returned for the last few seconds of the melody, and participants were then asked to report whether that last musical segment (probe) was correctly placed in time, considering the prior period of imagery (possible responses: yes; maybe yes; maybe no; no). Whereas half of these musical probes played the accurate continuation of the melody, the other half played an incorrect segment of the music (two beats early or late in the musical stream). For each melody, there were two types of probe start time: one probe was designed to start playing on a beat which was the closest to the 6 secs mark from the end of the melody *and* was the first beat in a bar (‘first’). The second probe started two beats earlier than the other probe, hence around the middle of the previous bar (‘middle’). Participants were tested on each melody 8 times overall: while tapping – ‘first’ played at the correct time, ‘first’ played at an incorrect time, ‘middle’ played at the correct time, ‘middle’ played at an incorrect time; These were repeated without tapping. The incorrect probes played the music starting +/− 2 beats from the correct start time. Each test round of a pair of melodies included two blocks for each of the movement instructions (i.e., w/ tap or w/o tap). Each block contained four trials. The order of the two melodies and the type of probe presented were randomized within each test round.

##### Pitch Test

Each trial in this test began with the first four seconds of the music, accompanied by the visual metronome. After the music stopped, the metronome carried on in silence, allowing participants to imagine the rest of the melody to the beat for about 4 or 8 seconds. Next, when the music had returned for the next few seconds of the melody, participants were asked to report whether the musical probe was played in the correct pitch, considering the prior period of imagery. The probes always played the appropriate part of the melody time-wise, but although half of them were played in the original pitch of the melody, the other half was shifted by a major third up or down. Pitch modification were implemented using librosa and preserved the melodic contour of the sequence. For each melody, there were 8 different trial start times that spread along the melody. The imagery lasted 4 sec for half of the trials (‘short’) and 8 for the other half (‘long’). The imagery end times (which are also a probe’s start times) were also spread along the melody. Participants were tested on each melody 64 times overall: while tapping – 4 different ‘short’, each one twice in a correct pitch and twice in incorrect, 4 different ‘long’, each one twice in a correct pitch and twice in incorrect; these were repeated without tapping. Each test round of a pair of melodies included four blocks for each of the movement instructions (i.e., w/ tap or w/o tap). Each block contained 8 trials. The order of the two melodies and the type of probe presented was randomized within each test round. One participant did not perform this test due to equipment malfunction.

For both Beat and Pitch tests, the order of the rounds (three melody-pairs) was randomized for each participant. After each trial, feedback was provided based on performance, in the shape of a green (correct) or red (incorrect) dot at the center of the screen. The blocks of the two movement instructions were interleaved, and half of the participants started their test with a block with instruction to tap. During the imagery periods, background noises recorded from an EPI sequence were played (in SNR of 0.25; implemented using librosa) to simulate the experimental environment in the scanner and to prevent participants from hearing other noises such as their tapping sounds and heartbeats. The auditory stimulus was played to participants using sound-proof headphones (Vic Firth Stereo Isolation Headphones). Responses were recorded using a portable silicon keyboard mounted on an Ester foam, in order to minimize potentially distracting tapping noises. The tests were run on Psychophysics toolbox (Brainard, 1997; Pelli, 1997; Kleiner et al., 2007) for MATLAB (MathWorks) in a sound-proof room. Using a microphone and an inspection window, an experimenter monitored for any potential sounds and movements produced by the participants.

Right-tailed *t* tests (*α* = 0.05) were conducted against chance level to assess the effect of memorization on accuracy of internal recall of melodies. Paired two-tailed *t* tests were conducted to compare the effect of tapping on accuracy of internal recall of melodies. Effect sizes were assessed using Cohen’s *d*.

#### Scanning Procedure

In the scanner, each participant performed the following conditions for each of the six melodies: (1) passively listened to the melody (“perception”); (2) silently imagined the music to the rhythm of a visual metronome – a bouncing ball at the center of the screen – while keeping motionless (“imagery w/o tap”); and (3) silently imagined the music to the rhythm of a visual metronome while tapping the right index finger to the beat of the music (“imagery w/ tap”). The order of the melodies was randomized across participants, and each melody run started with the perception condition, followed by the two types of imagery conditions, in a random order (Supp. Fig. 1).

Each imagery condition began with the first 6 seconds of the music (rounded to the nearest beat), accompanied by the same visual metronome as in the memory test (see *Learning phase*). After the music stopped, the metronome carried on in silence, allowing participants to imagine the rest of the melody to the beat. After each imagery task, participants were asked to report their level of confidence in the accuracy of their imagery and the level of the experienced vividness, using 1 to 7 scales. Runs in which participants reported confidence in their accuracy equal or lower than 5 were repeated, until confidence level was improved. The reported level of vividness was overall high across melodies and participants (w/o tap: M = 5.9, SD = 0.92; w/ tap: M = 6 SD = 1.06).

Participants also took part in control conditions, in which they watched the same visual metronome as in the imagery conditions while tapping to its beat (“control w/ tap”) or keeping motionless (“control w/o tap”), but were not asked to imagine the music. These two control conditions repeated twice: once in 73 BMP and another time in 107 BPM, controlling for the two tempo sub-groups of the melodies. The order of the control conditions was randomized across participants, with one BPM version at the beginning of the scan (i.e., before the perception and imagery conditions) and the other BPM version at the end of the scan.

Tapping and responses to questions were recorded using an MRI compatible two-button box in the right hand. Participants were provided with an MRI compatible in-ear mono earbuds (Sensimetrics model S14), which provided the same audio input in each ear. The volume of the auditory stimuli was adjusted individually for each participant to comfortable and clear level, and all the melodies were presented at the same volume level. Visual stimuli (metronome and self-report questions) were projected onto a rear-projection screen located behind the magnet bore, and were viewed with an angled mirror. Visual stimuli were created using the Psychophysics toolbox (Brainard, 1997; Pelli, 1997; Kleiner et al., 2007) and were combined with auditory stimuli using FFmpeg (http://www.ffmpeg.org/, version 4.0.2). The resulting audiovisual stimuli were presented and synchronized with MRI data acquisition onset using the Psychophysics toolbox.

During all runs, participants’ eyes were observed by an experimenter (using Eyelink 1000 eye tracker) to ensure they were engaged with the task and not falling asleep. In addition, the experimenter listened to the sounds coming from an MRI compatible microphone suspended approximately 1 inch from participant’s mouth, to ensure they were not vocalizing. Respiration was monitored during all runs at a sampling rate of 400 Hz using a belt fastened around the chest. The signal was transmitted from the scanner bed using a Seimens wireless device (respiratory sensor PERU).

### Data analysis

#### Preprocessing

The fMRI data were preprocessed in FSL (https://fsl.fmrib.ox.ac.uk), including slice time correction, motion correction, linear detrending, high-pass filtering (42s period, corresponds to the duration of the shortest melody), spatial smoothing (6 mm FWHM Gaussian kernel) and coregistration and affine transformation of the functional volumes to a template brain (MNI152). As a first step before preprocessing of the data, we cropped out the signals of the first 7-8 TRs of the beginning of each melody. This included the removal the period in which the music was playing at the beginning of the imagery conditions (4 or 5 TRs, depends on the tempo of the beat) as well as additional 3 TRs due to global arousal response after the end of the sound that added noise to the signal. All calculations were performed in volume space. Projections onto cortical surface for visualization were performed, as a final step, with NeuroElf (http://neuroelf.net). For voxel-wise analyses, functional images were resampled to 3 mm isotropic voxels.

#### Removal of respiratory signal sources

Slow changes in respiration over time have been shown to induce robust changes in the BOLD signal (Chang et al., 2009) in many cortical regions, especially in multiband sequences (Scheel et al., 2014; Golestani et al., 2018). Therefore, we used multiple linear regression to project out from the BOLD data the respiratory variation per time (RVT) nuisance variable and the RETROICOR correction algorithm. These calculations were modified to account for the interleaved slice order with simultaneous acquired slices (Scheel et al., 2014). The respiratory variables were projected out separately for each participant (except two who were missing a respiratory signal due to equipment malfunction) for all conditions before performing the preprocessing.

#### Inter-subject correlation analysis

We computed inter-subject correlation (ISC) in each condition for each of 400 parcels from independent whole-brain resting-state parcellation (Schaefer et al., 2018). ISC analysis maps were produced within each condition (i.e., motionless imagery, imagery while tapping, perception) of each melody, as well as between conditions (e.g., perception vs imagery) of the same melody and across different melodies. The ISC maps provide a measurement of the reliability of brain responses to each of the melodies in each of the conditions by quantifying the correlation of the time course of the BOLD activity across participants who share the same experience (Hasson et al., 2004; Hasson et al., 2009). For each parcel, ISC within a condition is calculated as an average correlation 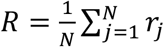, where the individual *r*_*j*_ are the Pearson correlations between that parcel’s BOLD time course in one individual and the average of that parcel’s BOLD time courses in the remaining individuals.

ISC across conditions is calculated as an average 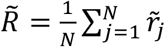 over the correlations, 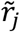 between the BOLD time courses of the *j*th individual from the first condition and the average BOLD time courses of all individuals in the other condition. When the comparison was performed across different melodies, the time courses of the longer melody of the two were cropped to fit the duration of the other melody, before the ISC was calculated.

When performed within a condition, the ISC method uses the participant’s brain responses within a given brain area as a model to predict brain responses to the experience shared by all.

When performed across conditions, the ISC method uses the averaged brain response to the experience in one condition as a model to predict brain responses to another experience in a different condition. The ISC was performed within each of the following conditions, for each of the six melodies: Imagery w/o tap, control w/o tap, perception. ISC was also performed across the following conditions: Imagery w/o tap versus perception, imagery w/ tap versus perception, control w/o tap versus perception, control w/ tap versus perception. Within each of these comparisons, ISC was calculate between both matching and non-matching melodies (or tempi) across the different conditions.

#### Inter-subject functional correlation analysis

We calculated the inter-subject functional correlation (ISFC) matrix between all parcels across brains of participants from 2 different conditions: perception versus imagery w/o tap, and perception versus imagery w/ tap. By measuring the inter-regional correlation across brains, instead of within a brain, the intrinsic neural correlations and non-neural confounds can be filtered out, thus increasing the ability to detect inter-regional coupling induced by shared experiences across participants (Simony et al., 2016). In addition, by also comparing the responses across conditions, we can identify couplings driven by the commonalities across perception and imagery, such as the internal representation of musical experience.

The neural signals *X*_*i*_ measured from participant *i*, *i* = 1, …, *k* are in form of a *p* × *n* matrix that contains signals from *p* neural sources (i.e., parcels) over *n* time points. All time courses were z-scored within participants to zero mean and unit variance. Thus, the participant-based ISFC was calculated by a Pearson correlation between single participant from one condition and the average of all participants from the other condition as follows: 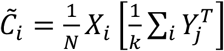, where the neural signals *Y*_*i*_ measured from participant *i*, *i* = 1, …, *k*. The cross-condition ISFC matrix was given by the following equation: 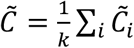. The final ISFC matrix is given 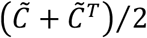, and then averaged with the final ISFC matrix calculated between individuals from the second condition and averaged signal from the first condition.

The ISFC was performed within each melody across the following conditions: Imagery w/o tap versus perception, imagery w/ tap versus perception, control w/o tap versus perception, and control w/ tap versus perception (Supp. Fig. 7).

#### ISC bootstrapping and phase-randomization

The statistical likelihood of each observed correlation was assessed using a bootstrapping procedure based on phase randomization, to preserve the long-range temporal autocorrelation in the BOLD signal (Zarahn et al., 1997). The null hypothesis was that the BOLD signal in each area in each individual at any point in time (i.e. that there was no ISC between any pair of participants). For all conditions, phase randomization of each parcel time course was performed by applying a fast Fourier transform to the signal, randomizing the phase of each Fourier component, and inverting the Fourier transformation. This procedure scrambles the phase of the BOLD time course but leaves its power spectrum intact. For each randomly phase-scrambled surrogate dataset, we computed the ISC for all areas in the exact same manner as the empirical cross-condition correlation maps described above. That is, for ISC within condition, the Pearson correlation was calculated between that parcel’s BOLD time course in one individual and the average of that parcel’s BOLD time courses of all other individuals. For ISC between conditions, the Pearson correlation was calculated between that parcel’s BOLD time course in one individual from one condition and the average of that parcel’s BOLD time courses of all individuals from the other condition. The resulting correlation values were averaged within each parcel across all participants, creating a null distribution of average correlation values for all parcels.

To correct for multiple comparisons, we selected the highest ISC value from the null distribution of all parcels in a given iteration. We repeated this bootstrap procedure 10,000 times to obtain a null distribution of the maximum noise correlation values for each melody (i.e., the chance level of receiving high correlation values across all parcels in each iteration), and then averaged each iteration value across all melodies to obtain a null distribution for the averaged ISC maps. Familywise error rate (FWER) was defined as the top 5% of the null distribution of the maximum correlations values exceeding a given threshold (*R*∗), which was used to threshold the veridical map (Nichols and Holmes, 2001). In other words, in the ISC map, only parcels with mean correlation values (*R*) above the threshold derived from the bootstrapping procedure (*R*∗) were considered significant after correction for multiple-comparisons and were marked as such on the final map.

#### Discriminability analysis

To test for the discriminability of the temporal responses to each of the melodies, we used a permutation analysis (Kriegeskorte et al., 2008) that compares the ISC between matching melodies against the nonmatching melodies (Baldassano et al., 2018). In each parcel in the brain, significant discriminability was determined by resampling correlations of non-matching melodies to generate a null distribution of the average across participants. That is, baseline correlations were calculated from nonmatching melody pairs, and compared to the average correlations of the matching melody pairs. This analysis ensures that the discovered temporal response patterns are content-specific at the melody level, as the correlation between neural patterns during matching melodies must on average exceed an equal-sized random draw of correlations between nonmatching melodies to be considered statistically significant. This procedure was performed for each parcel that was found significant in the ISC bootstrap analysis (see *ISC bootstrapping and phase-randomization*). The results were corrected for multiple comparisons across all significant parcels using False Discovery Rate (FDR; using *q* criterion of 0.05; (Benjamini and Hochbert, 1995)).

This discriminability analysis was performed on the ISC values calculated between the perception and imagery conditions, as well as on the ISC calculated within the perception condition.

#### Classification accuracy of individual melodies

We computed the level of discriminability of temporal patterns for individual melodies during perception, and across the perception and imagery conditions, in the parcels that showed significant ISC in the relevant condition. Participants were randomly assigned to one of two groups, an average time course was calculated within each group, and the data were extracted for each parcel of interest. When the classification was performed across conditions (e.g., perception and imagery), the data of each group were randomly sampled from one of the two conditions. Pairwise correlations were calculated between the two group means for all 6 melodies. For any given melody, the classification was labeled correct if the correlation with the matching melody in the other group was higher than the correlation with any other melody. Accuracy was then calculated as the proportion of melodies correctly identified out of 6 (chance level = 0.167). The entire procedure was repeated using 200 random combinations of the two groups sized *N* = 12 and *N* = 13. Statistical significance was assessed using permutation analysis in which for each combination of two groups, melody labels were randomized before computing the classification accuracy. Accuracy was then averaged across the 200 combinations and the procedure was performed 1000 times to generate null distributions for overall accuracy in a given parcel. Classification accuracy was corrected for multiple comparisons over all parcels of interest using FDR at threshold *q* = 0.05. The classification accuracy across the perception and imagery conditions was also calculated separately within each of the two tempo sub-groups of the 3 melodies (chance level = 0.334).

#### Inter-subject functional correlation enhancement by tapping

To measure the degree of neural response enhancement by tapping, we assessed the difference in reinstatement of the response to melodies during imagery when it was accompanied by tapping compared to motionless. Specifically, to measure effects of tapping on internal processing of melodies, in each brain area we subtracted the ISFC calculated between perception and motionless imagery, from the ISFC between perception and movement-accompanied imagery (i.e., ISFC(imagery w/ tap vs. perception) – ISFC(imagery w/o tap vs. perception)). Similarly, we also subtracted the ISFC calculated between perception and the control conditions (ISFC(control w/ tap vs. perception) – ISFC(control w/o tap vs. perception)). Significant enhancement of coupling across parcels was determined using a paired two-tailed permutation analysis that randomly assigned the correlations from the two imagery conditions (w/ or w/o tapping) to two new sham groups, generating a null distribution from the subtraction between the sham groups. We corrected for multiple comparisons across all parcels by controlling the FDR (*q* = 0.01).

## Code availability

Code supporting the findings of this study are available from the corresponding author upon request.

