## Supplementary Material for "Mapping the contents of consciousness during musical imagery"

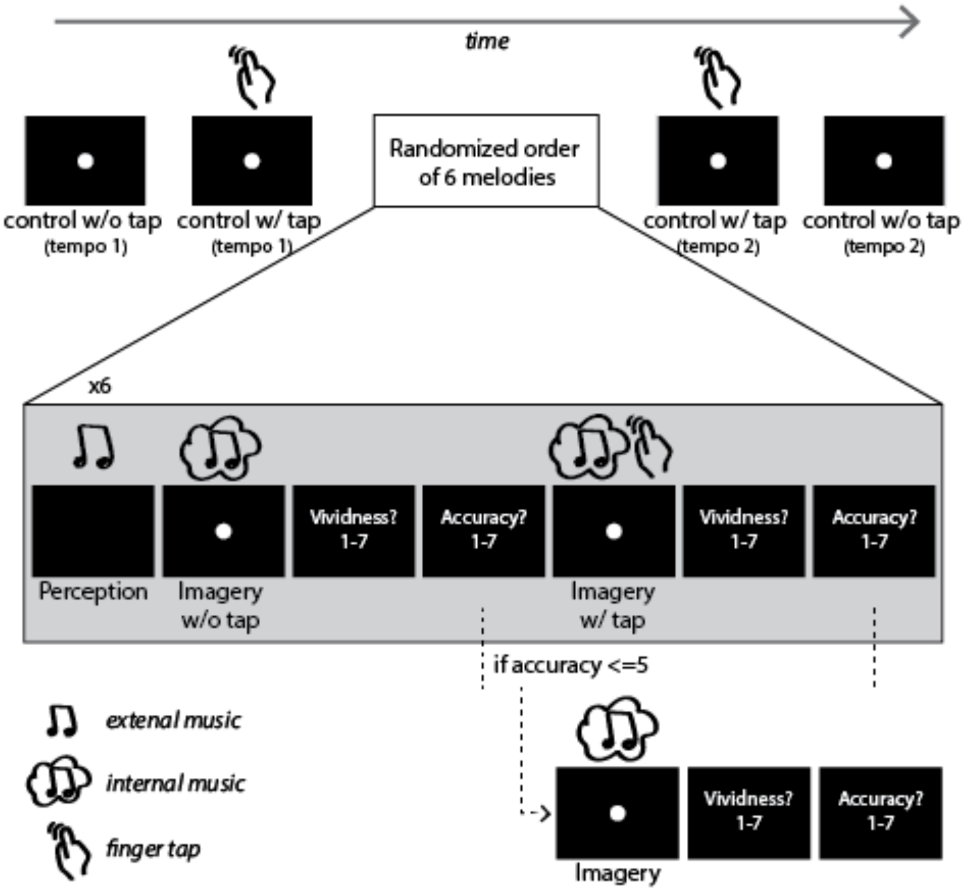


***Supp. Figure 1.*** *Schematic representation of the experimental design of an fMRI session. For each of the six melodies, participants passively listened to the melody ("perception") and then imagined it twice to the rhythm of a visual metronome (a bouncing white ball): while keeping motionless ("imagery w/o tap") and while tapping to the beat of the music ("imagery w/ tap"). After each imagery task, the level of confidence in accuracy of imagery and the level of experienced vividness was reported on a scale from 1 to 7. If the reported accuracy was equal or lower than 5, the imagery task repeated until confidence level was improved. At the start and end of the session, participants took part in control conditions, in which the visual metronome as presented, but they were not asked to imagine the music. The control condition was performed in the two tempi of the melodies (73 and 107 BMP), while tapping to the beat ("control w/ tap") or keeping motionless ("control w/o tap").*

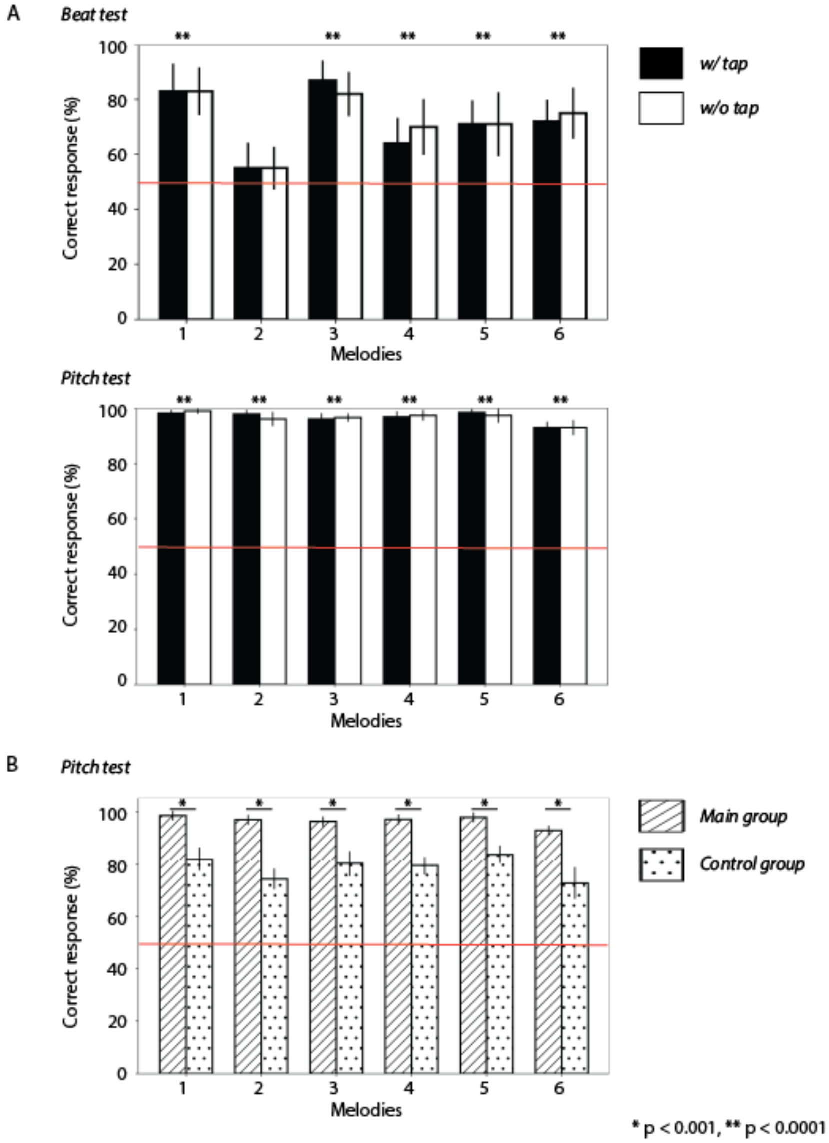


***Supp. Figure 2.*** *Imagery accuracy of the six melodies was assessed using Beat and Pitch tests, while either keeping still or tapping to the rhythm of a visual metronome.* ***A,*** *In both Beat and Pitch tests, designed to capture the temporal and tonal fidelity of the imagery, participants performed on average above chance (red line) for all melodies. Success rates were lower in the Beat than in the Pitch test, as previously observed in similar tests (Weir et al., 2015). These results, together with participants' ability to hum each melody out loud very accurately (see Methods, Experimental design), suggest that they were able to accurately imagine all the melodies. We did not observe an effect of tapping on accuracy of imagery in any of the tests.* ***B,*** *To test whether the extremely high performance in the Pitch test reflects the acquired knowledge about the melodies and not purely a general ability to track changes in musical key or register, we compared the performance of the experimental participants with an independent control group of 12 additional participants (8 females, ages 18-33, mean age 22.7) that performed the same Pitch test without memorizing the melodies in advance (see Supp. Methods, Experimental design). Participants who learned the melodies before the test had higher success rate in tracking the original pitch of the melody than the control group.*

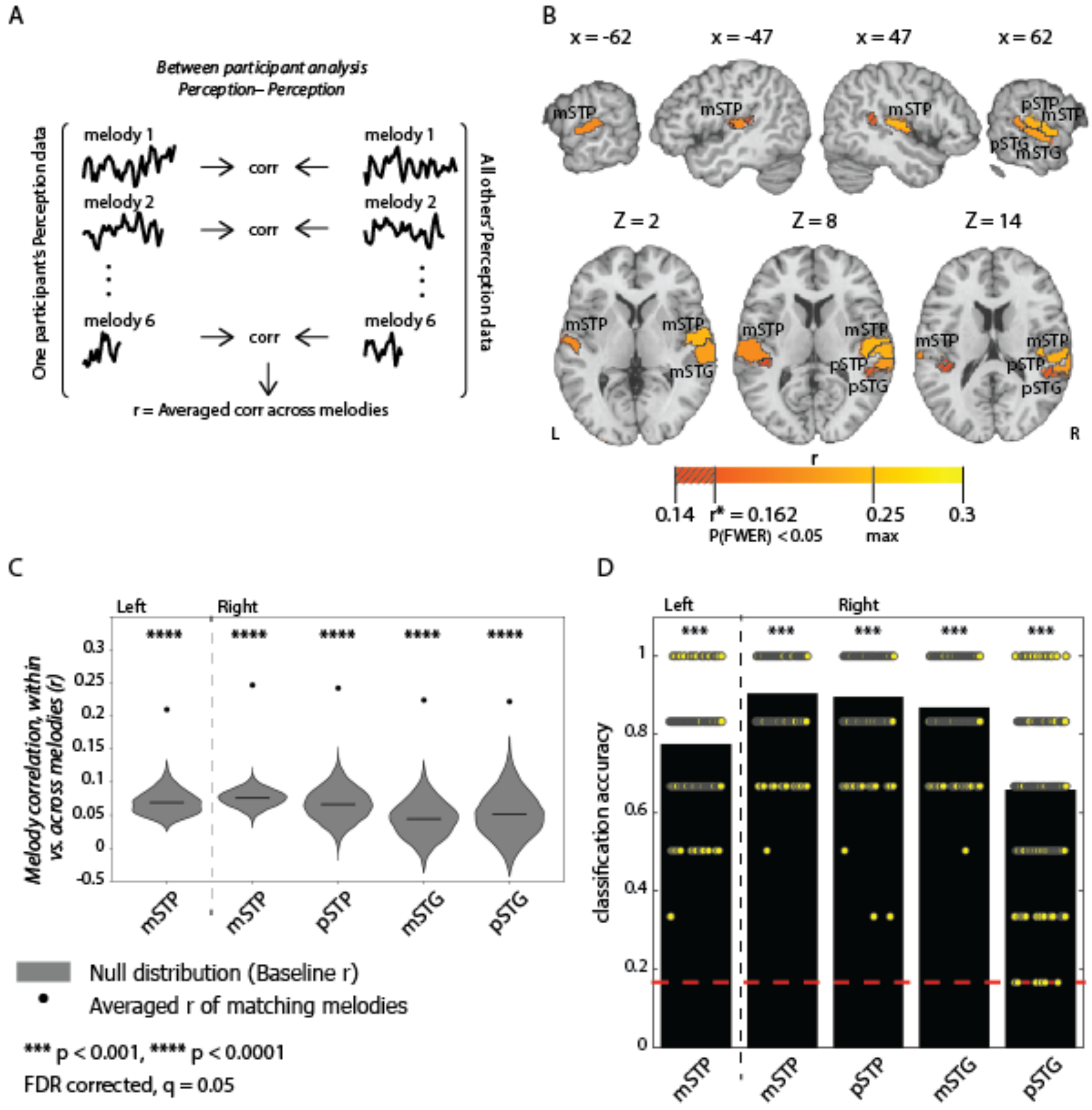


***Supp. Figure 3.*** *Temporal pattern similarity in the perception conditions.* ***A,*** *Schematic for between-participant analysis for the perception condition. BOLD time courses were correlated between participants (each participant versus the average of all others) for each of the six melodies in each parcel, to produce an averaged ISC during the imagery task.* ***B,*** *Cortical parcels where the highest temporal pattern similarity was observed across participants during perception of the same melodies ($p_{(FWER)} < 0.05$; threshold $r = 0.14$ for visualization purposes). Regions in the lateral STG include middle and posterior parcels (mSTG, pSTG); Regions in the STP include middle and posterior parcels (mSTP, pSTP).* ***C,*** *Discriminability of temporal patterns in five parcels that showed similarity during perception. Temporal patterns were discriminable from each other when the similarity between matching melodies was greater than between non-matching melodies. Black circles show average correlation of matching melodies. Statistical significance was determined by generating a null distribution of random*

pairs of non-matching melodies (grey violin baseline; FDR-corrected $q = 0.05$). ***D,*** *Classification accuracy of melodies across brains during perception. Participants were randomly assigned to one of two groups ($N = 12$ and $N = 13$), and an average time course calculated from the perception condition for all six melodies within each group. Accuracy was calculated as the proportion of melodies correctly identified out of six. The entire procedure was repeated using 200 random combinations of the two group sizes (yellow marks), and an overall average was calculated for each parcel (right mSTG 87%, pSTG 66%, mSTP 90% and pSTP 89%, and left mSTG 77%, black bars; chance level 16.7%, red line; FDR-corrected $q = 0.05$).*

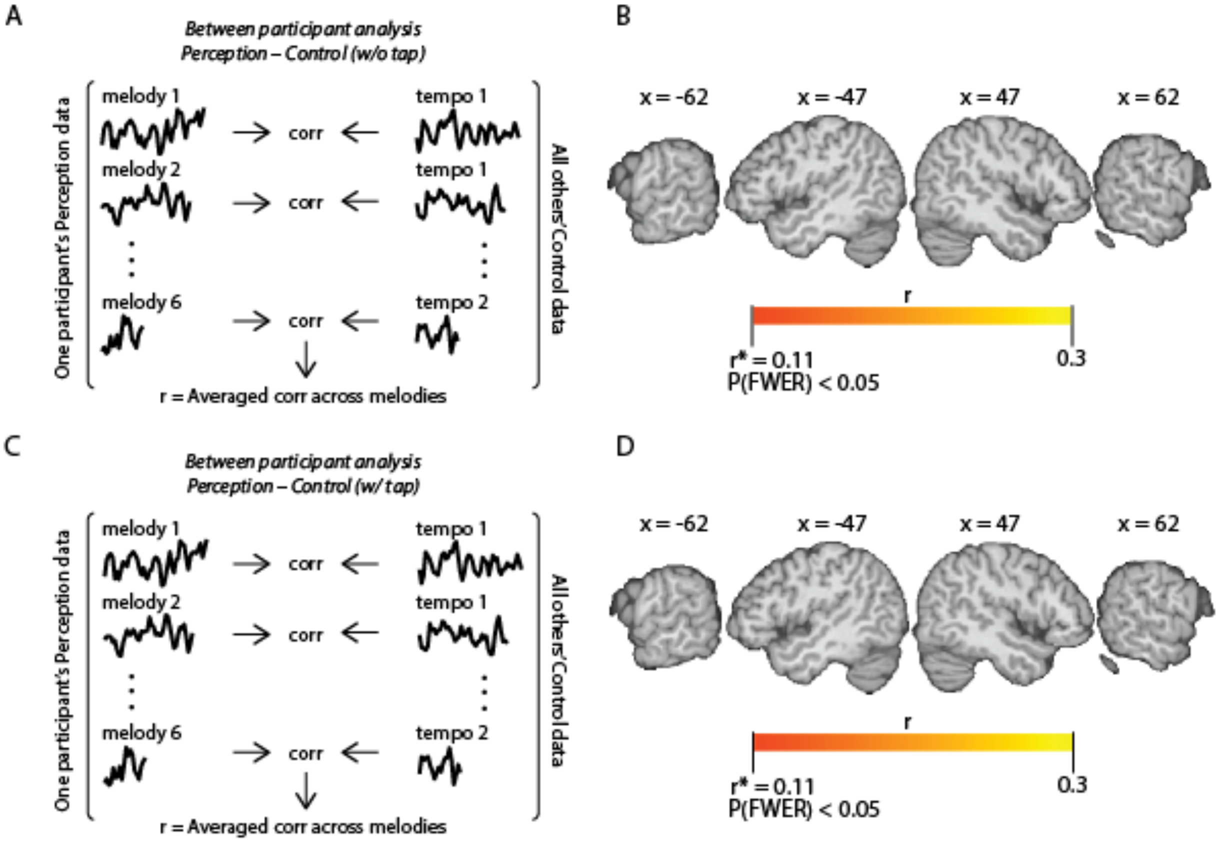


***Supp. Figure 4.*** *Temporal pattern similarity between control and perception.* ***A,*** *Schematic for between-participant similarity analysis across perception and control without tap. The correlations were computed between each melody during perception and its matching type of task tempi during the control (73 or 107 BPM).* ***B,*** *No cortical parcels in the temporal cortex show significantly similar temporal patterns between the perception and control (w/o tap) ($p_{(FWER)} < 0.05$).* ***C,*** *Same as A, except with tap.* ***D,*** *No cortical parcels in the temporal cortex show significantly similar temporal patterns between the perception and control (w/ tap) ($p_{(FWER)} < 0.05$).*

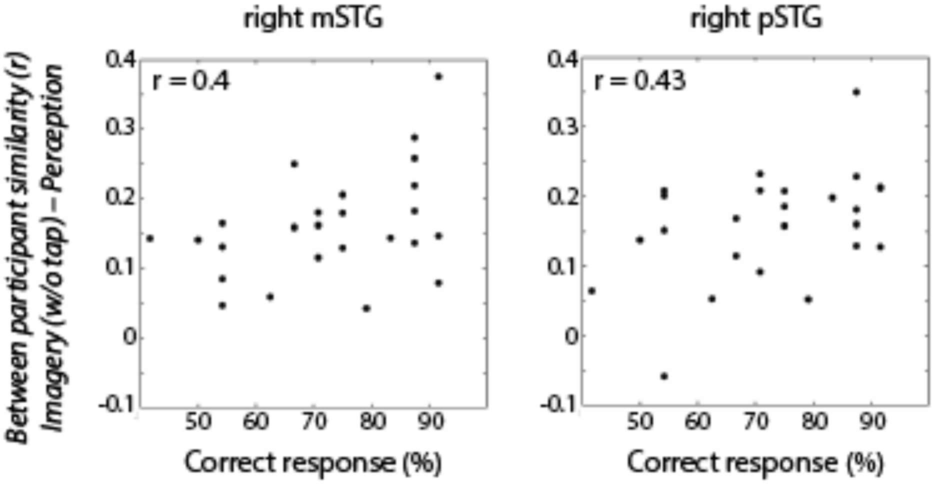


***Supp. Figure 5.*** *The greater the reinstatement of neural response during imagery (w/o tap) in the right middle and posterior STG, the better the imagery accuracy. We found a positive correlation between participant's strength of imagery-to-perception pattern similarity and their performance in the Beat test of imagery accuracy (right mSTG:* $r = 0.4$*,* $p = 0.047$*, pSTG:* $r = 0.43$*,* $p = 0.037$*, uncorrected for multiple comparisons).*

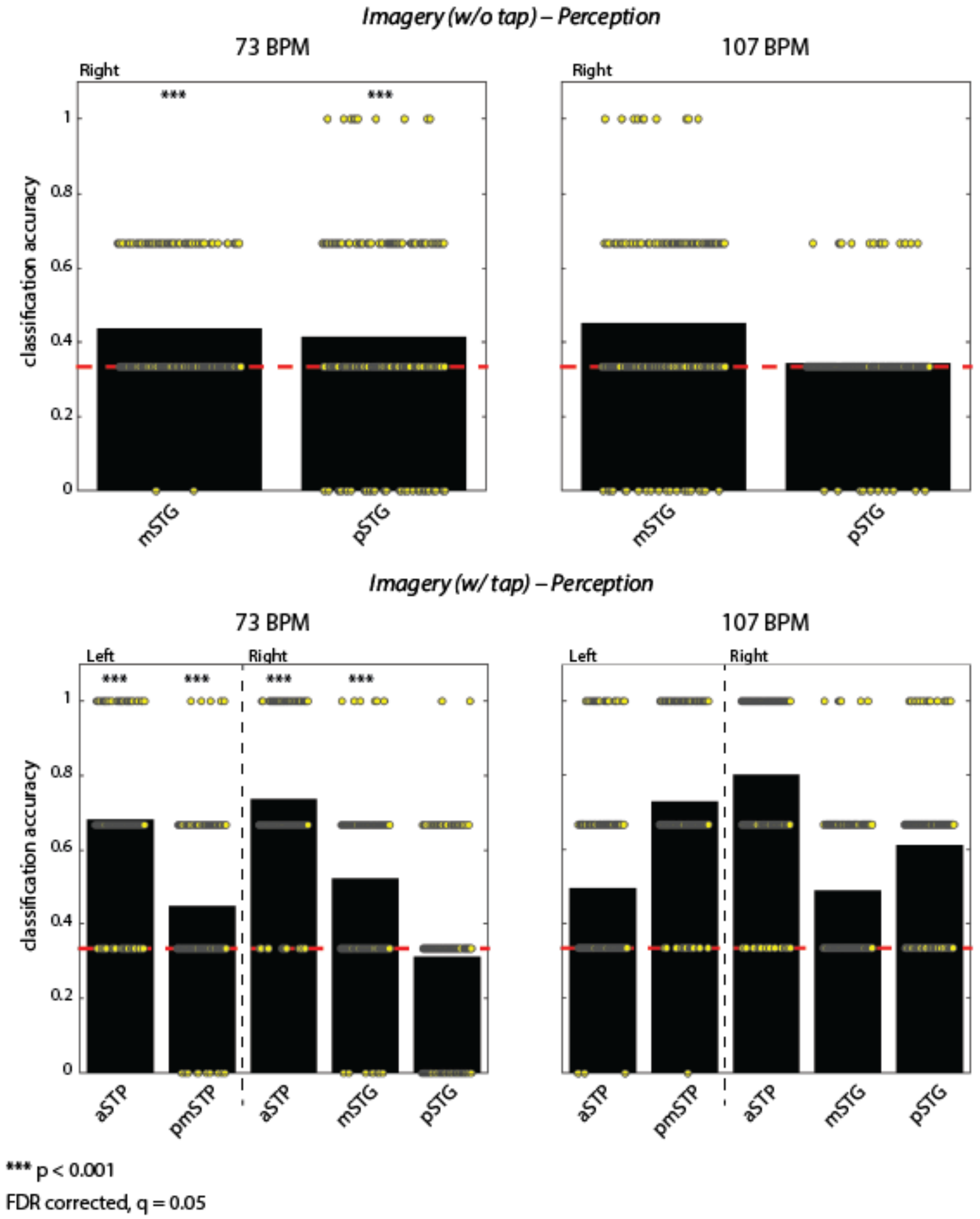


***Supp. Figure 6.*** *Classification accuracy of melodies across brains between imagery (w/ or w/o tap) and perception in the two tempo sub-groups of the melodies (73 and 107 BPM). Participants were randomly assigned to one of two groups (*$N = 12$ *and* $N = 13$*), and an average time course calculated from the imagery condition in one group and from the perception condition in the other for all three melodies in each tempo sub-group. Accuracy was calculated as the proportion of melodies correctly identified out of three. The entire procedure was repeated using 200 random combinations of the two group sizes (yellow marks), and an overall average was calculated for each parcel (black bars; chance level 33.4%, red line; FDR-corrected* $q = 0.05$*).*

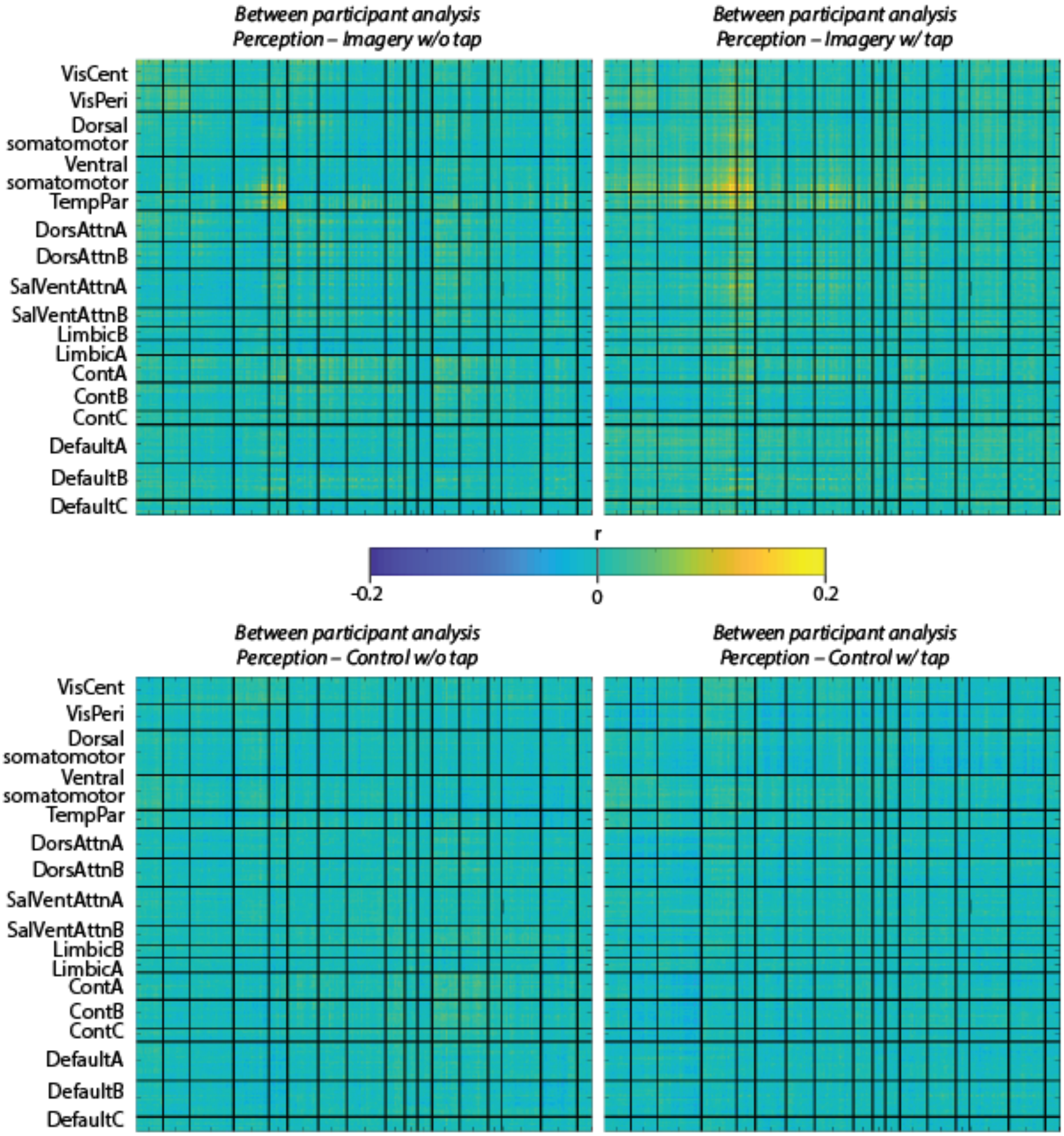


***Supp. Figure 7.*** *Inter-regional coupling in reinstatement of music during imagery and control (w/ and w/o tap). The inter-regional correlation of temporal responses was performed on a between-participant basis (see Methods, Inter-subject functional correlation analysis) for each melody between the perception and the imagery conditions across all parcels, which were organized in resting-state functional networks (Schaefer et al., 2018).*

**Supp. Table 1. MNI coordinates of parcels in auditory cortex (based on Schaefer et al. 2018)**

| | Area | R | A | S |
|---|---|---|---|---|
| Right hemisphere | mSTP | 54 | -14 | 6 |
| | aSTP | 52 | 4 | -6 |
| | pSTP | 60 | -24 | 10 |
| | mSTG | 62 | -18 | 0 |
| | pSTG | 64 | -34 | 10 |
| Left hemisphere | mSTP | -56 | -22 | 8 |
| | aSTP | -50 | -10 | 0 |
| | pmSTP | -40 | -36 | 14 |
| | pSTP | -58 | -36 | 16 |
| | aSTG | -52 | 6 | -12 |

### Supplementary Methods

*Behavioral assessment of imagery*

**Pitch Test – control group.** A separate group of participants (see Supp. Fig 2B) performed the Pitch test without any earlier exposure to the music. The test was identical to the one performed by the main group of participants who memorized the melodies. Since participants were not familiar with the melodies and could not be asked to imagine them, they were asked to try to determine whether the pitch of the music was shifted in the probe, compared to the few seconds of the melody played earlier.

To further explore the effect of memorization on the accuracy of the internal recall of melody's pitch, an independent sample two-tailed $t$ test was conducted between the main group of participants who learned the melodies and the control group who was not exposed to the melodies before the test.
